# RAStoERK: a reference interaction atlas for insights into signaling and disease

**DOI:** 10.64898/2026.04.07.716061

**Authors:** Georg Vucak, Sebastian Didusch, Leandro Cannizzaro, Ana Santiago, Markus Hartl, Jörg Menche, Manuela Baccarini

## Abstract

The RAS/RAF/MEK/ERK pathway has been extensively studied for its roles in physiology and disease, yet a systems-level view of its interplay with the broader cellular context is lacking, limiting insight into paralog-specific roles, mutation-driven rewiring, and cross-pathway integration. To address this, we simultaneously mapped the interactomes of all pathway components and their activating and inactivating mutations, identifying 2,500 high-confidence interactors, 88% of which previously unreported, and generating a high-confidence, comprehensive reference map of the pathway in resting and activated states. The pathway behaves like a “molecular organism” that connects to other signaling complexes, operates from distinct subcellular sites, and integrates into networks relevant to physiology and disease. Dataset exploration uncovered paralog- and mutation-specific interaction signatures and pathway-level crosstalk, including previously unknown connections to mRNA metabolism and WNT signaling. The interactomes available at *RAStoERK.univie.ac.at* provide a foundational community resource for unbiased discovery, targeted mechanistic studies, and potential therapeutic exploration.

**Highlights:**

- High-resolution interactome map of the RAS to ERK pathway in resting and active states
- State-dependent pathway-level interactions with signaling and functional modules
- Mutation-driven interactome rewiring and integration into pathophysiological networks
- A resource to discover pathway crosstalk, disease links, and actionable interventions

## Introduction

Cell physiology can be viewed as the product of thousands of proteins acting in concert to shape the cellular response to environmental signals. The transduction of these signals is accomplished by proteins arranged in cascades that transfer signals by way of posttranslational modifications, including phosphorylation/dephosphorylation. The correct timing and duration of the signal and its appropriate localization are essential for a productive outcome.

This coordination is achieved through protein-protein interactions (PPIs) that assemble functionally related proteins into signaling complexes, which are often tethered to organelles and ensure signal fidelity and strength. These signaling pathways are involved in extensive crosstalk, and generate interlocked, dynamic networks required to produce physiologically relevant outcomes. If any of these processes go awry, the whole organism may be at risk.

Indeed, PPIs have been shown to be perturbed in genetic disease.^1^ A case in point is the hyperactivation of the RAS/RAF/MEK/ERK signaling pathway caused by somatic mutations in RAS, the most potent human oncogene. RAS mutants are found in more than 20% of cancer cases, and these cancers have a worse prognosis than those lacking RAS mutations.^2,3^ RAS was long considered undruggable, but several strategies to target it have recently emerged, including allele-specific inhibitors or degraders. However, resistance mechanism including pathway reactivation or activation of parallel pathways complicate these therapies, and have led to the development of strategies co-targeting downstream effectors or other vulnerabilities.^4^

Downstream of RAS, the RAF/MEK/ERK kinase cascade is involved in the control of fundamental cellular processes as well as in the pathogenesis of human diseases. The RAS/RAF/MEK/ERK pathway comprises four tiers, each containing more than one enzyme (4 RAS, 3 RAF, 2 MEK, 2 ERK). RAF are also mutated in cancer, albeit with very different frequencies. The paralogs have distinct essential functions, some of which rely on PPIs and on their ability to crosstalk with other pathways rather than on their enzymatic activity. ^2,3,5,6^

Despite the extensive knowledge accumulated over the years, we still lack a comprehensive view of the interactions formed by pathway components and how they are perturbed by inactivating/activating mutations. Such insight is critical to predict the effects of degrader and combination therapies and to reveal pathway crosstalk. To address this gap and overcome these limitations, we used affinity purification (AP-MS) and TurboID-mass spectrometry (TbID-MS) to simultaneously mapped the pathway’s PPI network in natural and perturbed states (oncogenic or inactivating mutations), generating a pathway-wide reference set of “ground truth” interactions for future biomedical studies.

Our pathway-centric approach differs from efforts describing kinome-wide interactions, which, due to their global nature, sacrifice depth of analysis and exclude pathway components (e.g. in our case RAS). Studies focusing on one pathway component, on the other hand, are also limited because they only contribute to knowledge about individual proteins, not the pathway as a whole.

Interactome maps generated in different laboratories bear limited overlap, ^7,8,9,10,11,12,13,14,15,16^ possibly due to the use of different cell lines or different clones of the same cell line, different culture conditions, and/or different sets of controls. Therefore, interaction maps reconstructed by integrating datasets generated in different laboratories would be inherently unreliable. Our one-lab/one-pathway proteomic approach overcomes this limitation providing the most comprehensive description of the pathway interactomes to date.

## Results

### Building a Comprehensive Interaction Map of the pathway from RAS to ERK

To generate a pathway-level reference interaction map, we used one parental HEK293 cell line to produce stable cell lines containing Dox-inducible versions of the individual paralogs fused to V5-tagged BirA* (TbID; Figure 1A). We minimized the risk of identifying artifactual interactions caused by overexpression by titrating each bait to the expression levels of the corresponding endogenous protein. Streptavidin immunoblotting confirmed that all baits were capable of biotinylating proteins in vivo, and ERK phosphorylation was used to read out the activity of constitutively active (CA) or inactive (CI) upstream mutants (Figure S1A-B). CA-RAS and CA-RAF mutants represented the most common cancer-associated somatic mutations^17^ (updated from https://www.cbioportal.org/ querying curated sets of non-redundant studies; and from the cancer hotspots site https://www.cancerhotspots.org/#/home). The CA-RAS mutations disrupt GTP hydrolysis and guanine exchange rates,^3^ while the CI mutations increase the affinity of RAS for GDP.^18^ The two CA-BRAF mutations have distinct dependency on RAS and dimerization, while CA-RAF1 and, by extension, CA-ARAF mutations prevent the 14-3-3 binding that constraints activation.^3,5^ The CI-RAF mutations ablate the aspartate that chelates the Mg2+ ion that stabilizes the β- and γ- phosphate of ATP, and impact tumorigenesis^19,20^ or development.^21^

**Figure 1.**
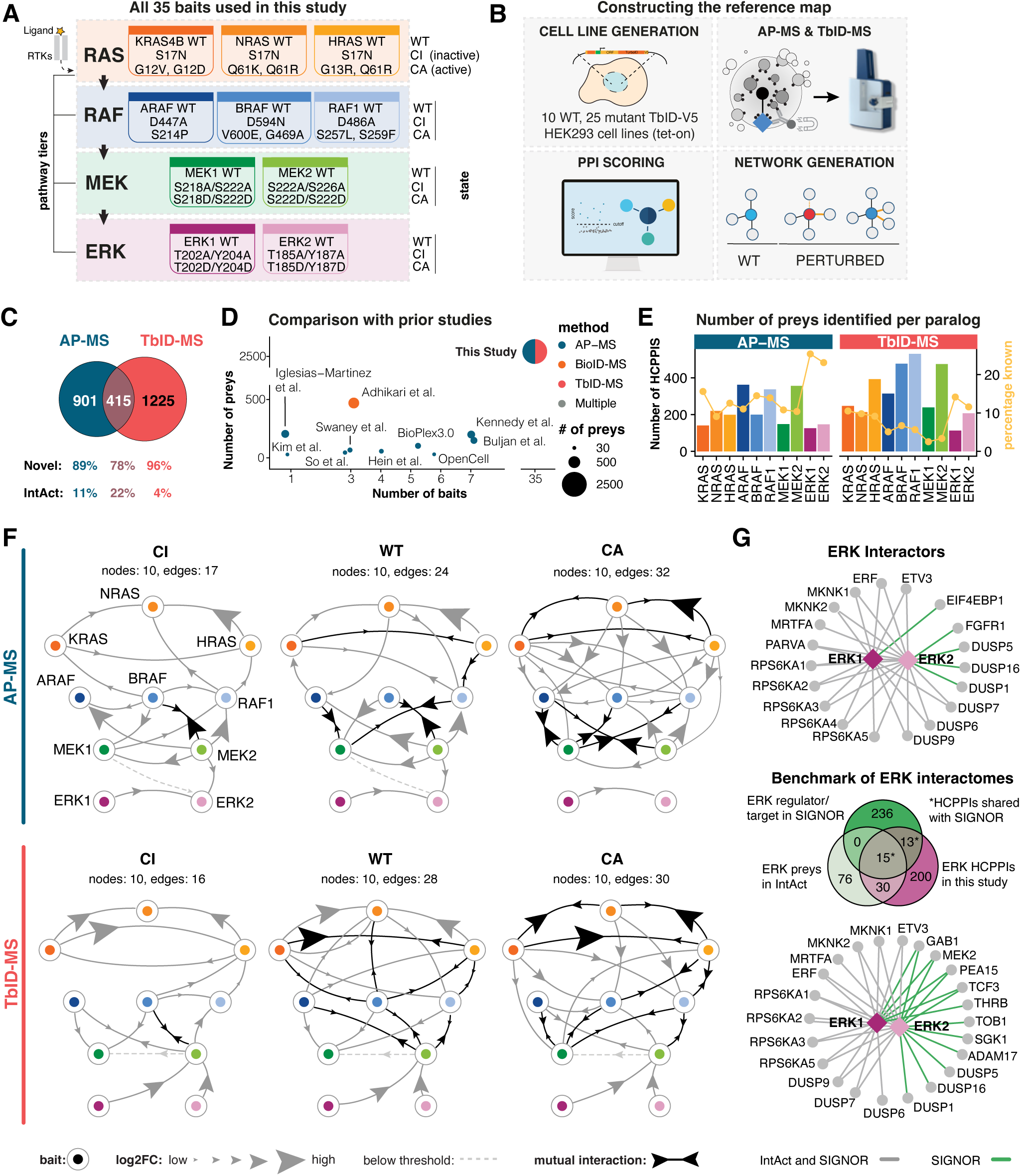
The largest library of ERK pathway interactions. (A) RAS/RAF/MEK/ERK pathway. Each of the 10 core proteins is represented by a different color and is mapped in 3 states (WT, CI, and CA), resulting in the 35 bait proteins depicted. The same color code is maintained throughout all figures. (B) Combinatorial interactomics screen workflow. See STAR Methods for details. (C) Number of HCPPIs detected by AP-MS and TbID-MS. The % of interactors listed or not in IntAct (novel PPIs) are shown. (D) Comparison with IntAct MS datasets using ERK pathway core proteins as baits. (E) Number of preys per bait. The bars are color-coded according to the bait and aggregated across CI, WT and CA paralog states. Yellow lines = percentages of known interactors. (F) Directed network of core pathway HCPPIs. The relationship between baits and preys is indicated by the direction of the arrowheads. Arrowhead size is proportional to log2FC compared to control. (G) ERK HCPPIs listed in SIGNOR and detected in this study. Green edges = HCPPI listed in <u>SIGNOR</u> alone; grey edges, HCPPIs listed in SIGNOR and IntAct. The Venn diagram shows the overlap among HCPPIs detected in this study, ERK interactors listed in IntAct, and causal relationships with ERK listed in SIGNOR.

MEK and ERK mutations in cancer are rare; however, their activation is defined by the phosphorylation of neighboring serine residues (MEK) or threonine/tyrosine residues (ERK). We thus used SS/DD or TY/DD mutations and SS/AA or TY/AA mutations of the activation sites to mimic activation or prevent it. Interactors isolated via V5 affinity purification (AP) or streptavidin pulldown from biotinylated cell lysates (TbID) were identified by diaPASEF-MS,^22^ resulting in deep interactome coverage and accurate quantification (Figure 1B).

Our study features the largest number of RAS/ERK pathway baits (35 between WT and mutated) reported in any previous high-throughput datasets,^7,8,9,10,11,12,13,14,15,16^ and identifies the largest number of high-confidence PPIs (HCPPIs), defined as proteins significantly enriched over the GFP control with a log_2_FC threshold equal to, or greater than that of known interactors. AP-MS and TbID-MS generated highly complementary interactomes, with a 16% overlap consistent with that reported in other studies employing both technologies.^23,24^ The two methods combined identified more than 2500 HCPPIs, a substantial improvement on previous high-throughput studies. For comparison, the second largest studies used 7 baits each and identified a maximum of 200 interactions. 88% of the HCPPIs identified in our study were not present in the IntAct^25^ database (Figures 1C-D; Table S1). RAF and MEK baits had the largest interactomes with the highest percentage of previously undiscovered interactions (Figure 1E).

### Quality control and first insight into pathway architecture

Principle component analysis (PCA) on the bait – prey intensity matrices showed tight clustering of triplicate samples, indicating low biological and technical variability in the datasets. PCA (Figure S1C) and hierarchical clustering of the overlap coefficient for all pairwise comparisons between sets of interactors (Figure S1D) provided an initial view of pathway architecture. The interactomes largely clustered along the four pathway tiers, as expected given the high bait homology in each tier. RAS and ERK tiers clustered away from RAF and MEK, and within the RAS tier, KRAS interactomes separated from H and NRAS. Across tiers, the largest overlap was observed between RAF and MEK, with CA BRAF interactomes closer to the MEK interactomes than to those of ARAF and RAF1 in the AP-MS dataset. This pattern was not observed in TbID datasets, where all CA-RAF and CA-MEK interactomes clustered near one another, albeit less tightly.

### Benchmarking, Coverage, and Discovery

The components of the RAS/RAF/MEK/ERK pathway are interconnected through PPIs. RAF^26,27^, MEK,^28,29^ and ERK^30^ paralogs heterodimerize within their own tiers. In addition, physical cross-tier interactions (e.g. RAS/RAF; RAF/MEK; MEK/ERK) occur upon activation.^5,6,31,32^ We benchmarked our datasets against these known interactions and assembled directed PPI networks of the core pathway. Nodes represent baits and endogenous preys; arrows point from bait to prey. For clarity, 14-3-3 and other pathway-related proteins such as KSR were omitted (Figure 1G). AP-MS and TbID-MS networks were broadly similar, though not identical, and are discussed together. In WT networks, RAS interacted with paralogs within its tier, with HRAS favoring NRAS. We also observed interaction between MEK2 and MEK1, and ERK1 and ERK2, but not among the RAF paralogs. Across tiers, the RAF/MEK module emerged as the pathway’s cornerstone. RAF connected to at least one RAS, with KRAS preferentially capturing RAF1 and BRAF. All three RAF captured MEK, but MEK2 engaged more than MEK1, both with RAF upstream and within its tier. Finally, we find MEK and ERK connected by significant albeit mostly low-strength interactions, and ERK baits connected mostly to MEK2.

CI and CA mutations induced opposite connectivity changes: CI reduced both the number and strength of cross-tier connections, effectively disconnecting CI-RAS baits from RAF. By contrast, CA increased connectivity, with most CA-RAS linked to RAF. RAF paralogs now connected with one another, faithfully recapitulating the activation-dependent heterodimerization described in the literature, and all RAF connected to MEK.

Our combinatorial approach extends prior work that used all RAF/MEK/ERK baits, by recovering connections extensively described in low-throughput studies (RAS paralog interactions, HRAS interactions with ARAF and RAF1, and cross-tier MEK–ERK links; Figure S1E), and by faithfully capturing pathway rewiring in resting and activated states.

Compared with the other tiers of the pathway, ERK tier interactomes were smaller in size and contained a comparably high number of known interactors and therefore represent a conservative benchmark for coverage and discovery potential against prior MS-based datasets (IntAct). Our interactomes contained more ERK HCPPIs than IntAct and captured 37% of those reported there, including substrates (RPS6Ks, MKNKs) and DUSPs that dephosphorylate ERK (DUSP7, DUSP9) or are regulated by ERK2 (DUSP16). Twenty-eight HCPPIs were indexed in <u>SIGNOR</u>^33^ meaning they have experimentally validated causal relationships with ERK (activation, inhibition, phosphorylation). These encompassed all 15 IntAct ERK preys indexed in SIGNOR and almost as many (13) additional HCPPI listed in SIGNOR but not in IntAct (Figure 1G).

Thus, our focused, pathway-wide interactome approach confirms the results of prior high-throughput studies but outperforms terms by recovering the highest number of preys, achieving an unprecedented degree of connectivity among pathway components as well as between ERK and its HCPPIs, and recapitulating perturbation-induced pathway rewiring.

### State-specific interactome remodeling

To understand the extent and structure of interactome dynamics beyond the rewiring of the pathway itself, we analyzed the impact of CI/CA perturbations on the distribution of HCPPI within pathway tiers (Figure S2A-C). WT RAF and RAS captured the highest number of preys, and capture was increased by both CI and CA mutations. WT-MEK retrieved few preys and showed the highest HCPPI gain, particularly in the CA condition. Prey capture by ERK, in contrast, was highest in the WT condition, and both the CI and the CA mutants decreased it. These results were consistent in both the AP-MS and TbID-MS datasets, although TbID contributed more HCPPIs to the combined dataset.

Diving deeper, we analyzed how CI/CA mutations impacted paralog-specific interactomes (Figure 2A-B). Interactome size varied greatly among paralogs of the same tier, and for RAS and RAF, the evolutionarily younger paralogs KRAS, NRAS^34^ and ARAF,^35,36^ showed the highest number of interactions in the WT state. This diversification of interactions may indicate distinct biological roles for different paralogs, consistent with *in vivo* experiments involving specific pathway members.^2,35^ Additionally, we detected both state-agnostic and state-specific HCPPIs, the most striking example of the latter being the substantial increase in interactome size caused by CA mutations in all baits except ERK. Typically, a majority of the HCPPIs common to multiple paralogs were new, CA-specific interactions, but some redistribution of HCPPIs from CI and/or WT interactomes of individual paralogs into common or shared CA interactomes was also observed. Examples of this are the RAS preys found in the stream containing RAF, or the RAF preys found in the streams containing HRAS or MEK2, and the MEK preys in the streams containing RAF. Notably, CA-ERK interactomes showed a decrease in common preys. This indicates that the HCPPIs—and possibly the functions—of paralogs in the upper tiers converge upon activation, consistent with the reports on RAS^37^ and RAF^26,27^ oligomerization, whereas in the lower tiers they rather diverge (Figure 2B).

**Figure 2.**
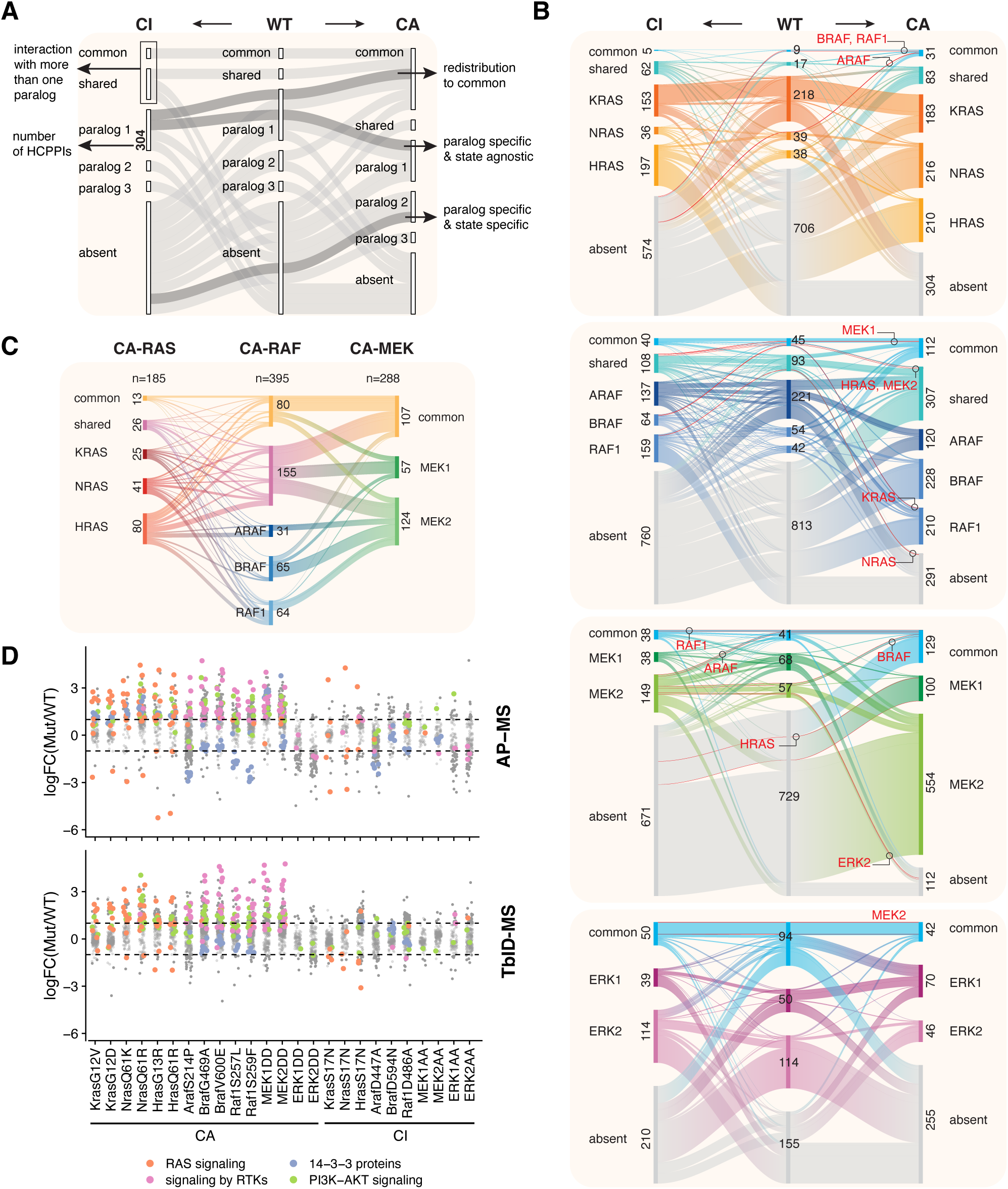
CI and CA mutations impact the flow of information through the ERK pathway. (A) Visual legend explaining the patterns displayed in the alluvial plots. (B) Alluvial plots represent the (re)distribution of interactors across the CI, WT, and CA states of individual paralogs, shared by two paralogs or common to all paralogs in a tier. The width of the bands is proportional to the number of interactors, shown next to the nodes. Streams containing pathway components are shown in red. In the case of a protein having two activating mutants, interactomes were combined into one entity per state. (C) Alluvial plot showing the flow of HCPPIs across tiers in CA interactomes. (D) Quantitative comparison of paralog- and mutant-specific interactomes. Log2FC larger than +1 or -1 (mutant/WT) were defined as GOI or LOI, respectively. Each dot represents an HCPPI. Components of selected signaling pathways are in color.

CI/CA mutations also impacted HCPPI overlap across tiers. Overlap was generally low in WT state, except for the MEK sample (55%; Figure S2A). CI mutations reduced the number of HCPPI shared among RAS/RAF/MEK and between the MEK/ERK tiers. Conversely, tier overlap clearly increased in the CA baits. ERK was again the exception, sharing very few HCPPIs with other tiers. Notably, overlap across tiers was more prominent in the TbID set (Figure S2B), which was thus overrepresented in the combined MS set.

Following RAS activation, pathway components engage in intra-pathway interactions and in crosstalk with other signaling systems, and this has informed the design of vertical and horizontal combinatorial therapies.^4^ Consistent with this, 25% of the RAS interactomes were shared with RAF paralogs, with flows from HRAS dominating the overlap. RAS interactomes overlapped preferentially with the HCPPIs of RAF complexes (common or shared), which comprised 60% of all RAF PPIs interacting with RAS or MEK. BRAF- and RAF1-specific interactomes contributed equally to the overlap with RAS and MEK, while less ARAF preys were shared with the other tiers of the pathway. Downstream, the MEK2 interactome and the common MEK1/MEK2 interactomes contributed mostly to the overlap with the other tiers (Figure 2C).

Consistent with the increased interactome overlap in the CA-state at the overall pathway level, analysis of fold changes in interactor intensity revealed a strong gain of interaction (GOI) in HCPPIs involved in RAS, RTK, and PI3K signaling for all individual CA-baits. Loss of interaction (LOI) was also detected. The most evident LOI, observed in the AP-MS dataset, was that of the 14-3-3 proteins with the ARAF and RAF1 mutants ablating one of the 14-3-3 docking sites, again validating the results of our screens (Figure 2D).

Notably, all CA interactomes contained unique HCPPIs that did not overlap with the other tiers, ranging from 75 % of the CA RAS interactome, to 60% and 64% of the CA RAF and CA MEK interactome, respectively.

Thus, constitutive activation increased both the interaction potential of all pathway components and the overlap across pathway tiers, at least partially by pathway-level engagement with other signaling systems. The substantial fraction of interactors unique to specific nodes indicates a vast potential for the discovery of functional paralog-specific interactions.

### Extensive pathway-level interactions with other signaling modules

In this context, mapping our interactomes on the kinome tree^38,39^ revealed interactions with about 19.5% of the kinases, confirming known partners (e.g. KSR for RAF and MEK, COT and PAK for MEK, RSK for ERK)^5,6^ and revealing new ones. Examples are CLKs, potential drug targets regulating splicing^40^ found in the RAF interactomes; or AURKA, PLK1, BUB1 and BUBR1, all involved in mitotic progression^41^ found in the MEK1-specific interactomes. The MEK2 TbID interactome contained CDC25C, a phosphatase activated by, and found in complexes with PLK1,^42^ reinforcing the connection between mitotic signaling and MEK (Figure 3A-B and S3A-B).

**Figure 3.**
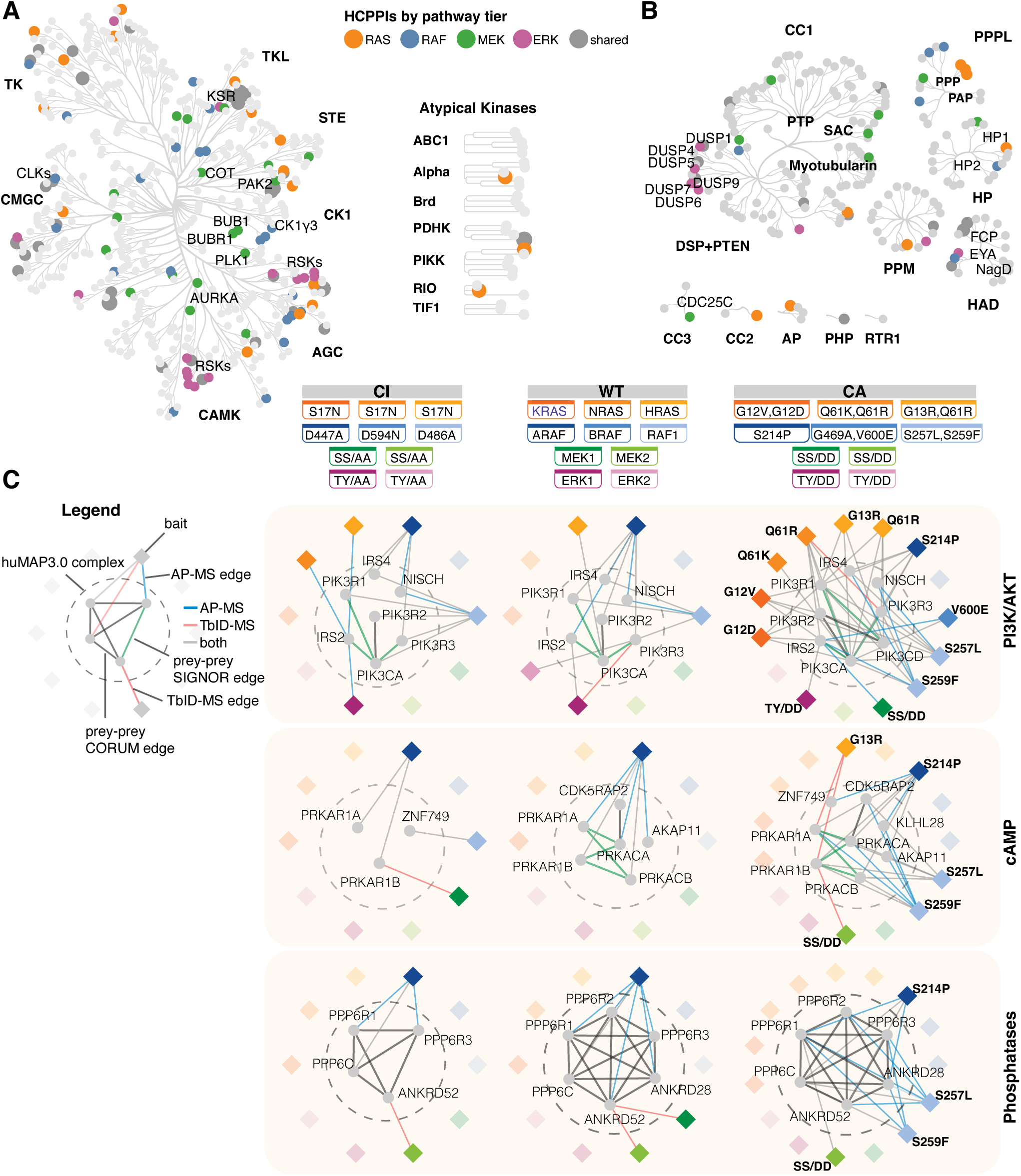
Pathway interaction with signaling complexes. (A, B) AP-MS/TbID-MS interactomes mapped on the kinome/phosphatome evolutionary trees. HCPPIs are color-coded by pathway tier, aggregated across states. Kinase/phosphatase families are bold. Shared = proteins interacting with multiple tiers. (C) Pathway interactions with representative huMAP3.0 complexes. Bait-prey edges are color-coded by detection method; high-confidence interactions among preys are in black (CORUM) or green (SIGNOR). Baits that interact with the complexes are highlighted.

Similarly, about 20% of the phosphatome^43,44^ was represented in our interactomes. Prominent HCPPIs were PPPLs connected mostly with RAS and RAF, and dual-specificity phosphatases^45^ (DUSPs) targeting ERK. The DUSPs were primarily detected by the ERK baits, but some interaction was observed between activated BRAF, RAF1 and MEK with DUSP1 and 5. The cytosolic, ERK-specific DUSP6 and 7 interacted only with the ERK baits, highlighting the selectivity of our system (Figure 3A-B and S3A-B).

Kinases and phosphatases often signal in the context of complexes that define their specific function. To assess whether our baits engaged with individual enzymes or rather with signaling complexes, we combined information from the hu.MAP3.0 resource,^46^ which integrates complexes identified by multiple approaches, from the CORUM database of experimentally validated mammalian protein complexes,^47^ and from the SIGNOR database,^33^ which annotates causal interactions not limited to physical complexes. We identified several complex interactions, in which connectivity increased with the activation state of the baits (Figure 3C). For example, HRAS, ARAF, RAF1 and ERK1 interacted with PI3K signaling complexes^48^ independently of activation state, while other RAS/RAF paralogs and MEK1 connected only in their CA state. Paralog preference was evident in the cAMP^49^ example, where ARAF interacted with at least one PKA subunit irrespectively of its state, while other pathway components (RAF1, MEK2) only connected in the CA state. A similar pattern was seen with PP6^50^, where ARAF and MEK2 engaged in all three states and RAF1 only in the CA state.

Importantly, paralogs also connected to subunits targeting holoenzymes to specific substrates, providing context for future functional studies. Examples are the interaction of ARAF and RAF1 with AKAP11,^51^ a LIR-containing scaffold connecting PKA with the autophagic machinery; or the connection to ANKRD52,^52^ which targets PP6C to PAK1 and is implicated in cell motility and metastases; and to ANKR28,^53^ which targets PP6 to focal adhesions to regulate their dynamics during motility (Figure 3C). Thus, ERK pathway components, particularly in the CA state, coordinately engaged multiple elements of other signaling complexes, generating complex networks of pathway-level interactions that are only detectable within a complete, pathway-profiled dataset and that provide crucial context for future functional studies.

### Subcellular, state-specific landscape of pathway interactions with functional networks

ERK pathway components signal from different subcellular locations, to which they redistribute during activation and/or to which they are tethered by scaffold proteins promoting their interaction with specific substrates, and ultimately determining different outputs. Classic examples are membrane recruitment of RAF by activated RAS, or phosphorylation-dependent nuclear translocation of ERK.^5,6,54^ We therefore used the subcellar UMAP provided by <u>organelleIP</u>^55^ to map the subcellular distribution of AP- and TbID-MS interactomes (Table S1).

CI and WT interactomes tended to co-localize in both datasets, while CA interactomes showed tier- and paralog-specific subcellular redistribution, particularly in the TbID dataset (Figure S4). For instance, many KRAS HCPPIs mapped to the ER, while multiple HRAS interactors mapped to plasma membrane and actin cytoskeleton; in the RAF tier, RAF1, BRAF and, to a lesser extent, ARAF preys also mapped to the plasma membrane. In addition, BRAF and RAF1 interactors were overrepresented in the ER. MEK HCPPI mapped to both ER and stress granules, while ERK interactomes were somewhat enriched in the nuclei. Most of these changes are consistent with the established relocalization patterns of active pathway components (Figure S4A, Table S1).^5,6^

Pathway enrichment analysis performed on the complete interactomes was not informative, yielding general (“signaling”) or pathway-related terms. In contrast, interactome analysis stratified by subcellular location returned more specific terms, showing enrichment in crucial biological processes, some of which previously associated with pathway components. Many terms were common to several paralogs in a tier, but paralog specificity was also observed (Figure 4A). Figure 4B-E shows examples of biologically relevant Reactome^56^ complexes enriched in specific subcellular localizations, aggregated across states. As expected, the TbID dataset contributed most interactions, but a core set was common to both methods, and AP-MS contributed preys critical for the completion of subcomplexes.

**Figure 4.**
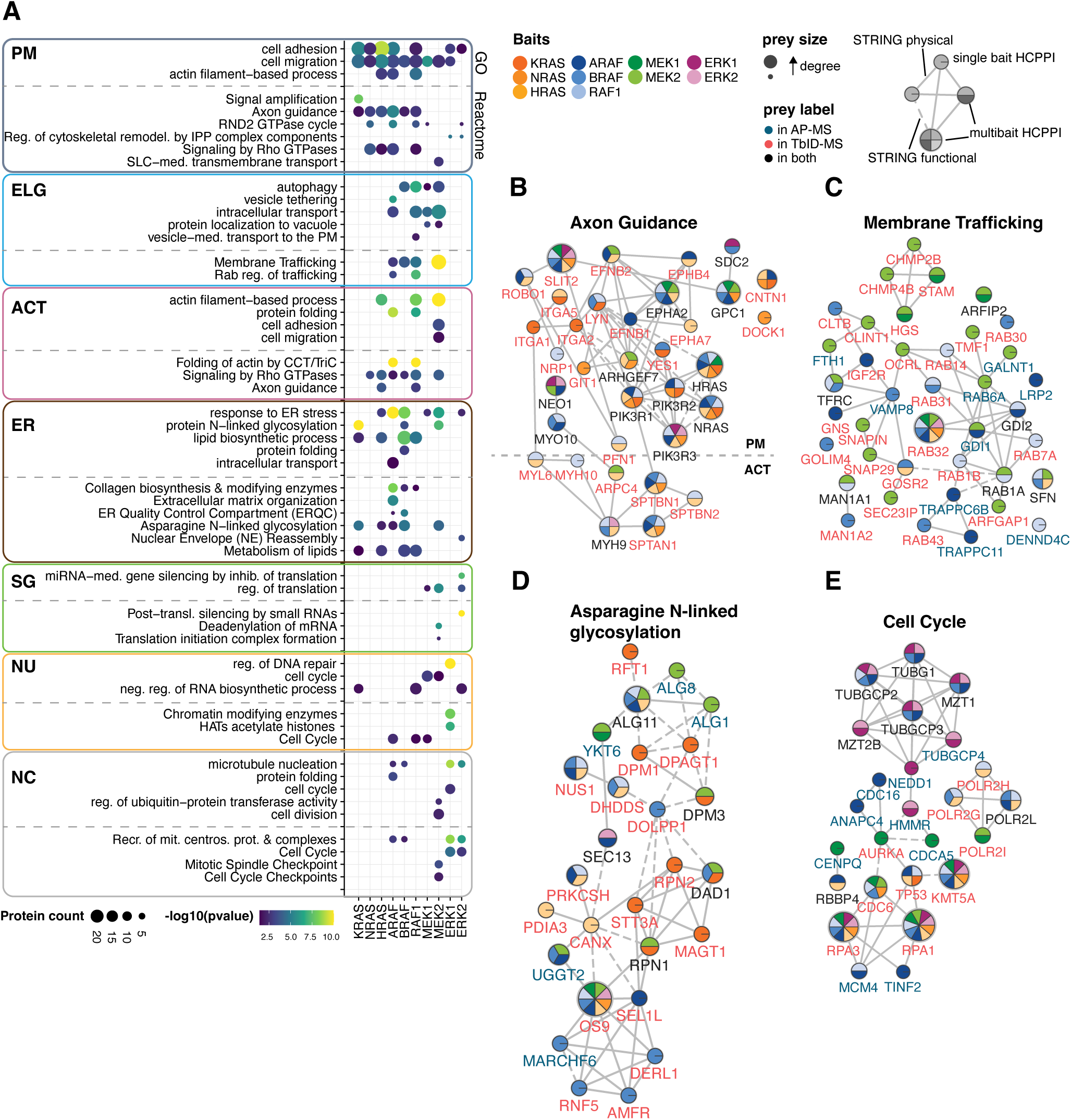
Subcellular landscape of interactions. (A) GO and Reactome enrichment analysis of combined AP-MS and TbID-MS interactomes aggregated across CI, WT and CA paralog states and stratified by subcellular localization according to <u>organelleIP</u>. In the dot plot, dot size is proportional to the number of proteins in each term and color reflects -log10(adj. p-value). PM = plasma membrane; E/L/G = Endosome/Lysosome/Golgi, combined endomembrane organelles; ACT = Actin cytoskeleton; ER = endoplasmatic reticulum; SG = stress granules; NU=nucleus; NC = not classified. (B-E) STRING networks of HCPPIs involved in selected Reactome pathways. Nodes are colored according to the paralogs they interact with. Node size is proportional to the number of interactions with the baits (node degree). Blue labels = nodes detected by AP-MS; red = detected by TbID; black = detected by both. Solid edges = STRING physical interactions (minimum score 0.4); dashed edges = STRING high-confidence functional interactions (minimum score 0.75).

The axon guidance complex localized at the plasma membrane/actin cytoskeleton contained many shared nodes (larger, multicolored nodes), and HRAS, ARAF, and MEK2 were strongly represented baits. In contrast, membrane trafficking and N-linked glycosylation complexes contained more bait-specific nodes. KRAS, BRAF and MEK2 were strongly represented in the N-linked glycosylation complex, while RAF, particularly RAF1, and MEK2 interacted strongly with the components of the membrane trafficking complex. ERK were prominent interactors of the cell cycle complex, mostly localized in the nucleus.

Analysis of state-specific networks (Figure S4B) revealed activation-induced expansion, with CA most closely matching the aggregated sets. The effect was strongest in axon guidance and weakest in cell cycle. Although the topology of the four networks differed, CA consistently recruited peripheral machinery—receptors, cytoskeletal components, ESCRT, ER-QC/ERAD, and APC/cohesin—around cores conserved across all three states. Many of these peripheral nodes were engaged by specific paralogs.

These results show that the interactomes in this study achieve remarkable resolution at the paralog level, support the notion that specific members of the pathway engage in concerted interactions with functional and/or physical complexes at defined subcellular localizations, and highlight that, together, the AP and TbID-MS datasets provide complementary, functionally relevant information.

### Mapping disease-relevant interactions

Given the clinical relevance of the pathway, we analyzed the distribution of high confidence disease-associated proteins (HCDPs), defined using the DisGeNET v25.2 database,^57^ across the interactomes (Table S1). HCDPs numbers increased in CA interactomes, ranging from negligible (KRAS) to extremely strong increases (MEK2; Figure S5A). As noted before, CA AP-MS and TbID networks were similar but not identical. For instance, SHOC2-linked phosphatases (PP-1A, B, and G)^58^ or the TRiC complex^59^ were present in the CA-AP-MS network; by contrast, the ARHGEF7-GIT1 complex^60^, AKT3, and the TNRC6A, B, C proteins,^61^ required for miRNA-dependent repression of translation, were selectively enriched in the CA-TbID-MS network (Figure S5B, C). In all networks, we observed paralog-specific HCDPs (most evident for NRAS Q61R and MEK2), tier-specific HCDPs (e.g. in the RAS tier) and tier overlapping HCDPs, visibly concentrated at the intersection between CA-RAF and CA-MEK tiers (Figure S5B-C; Figure 5).

**Figure 5.**
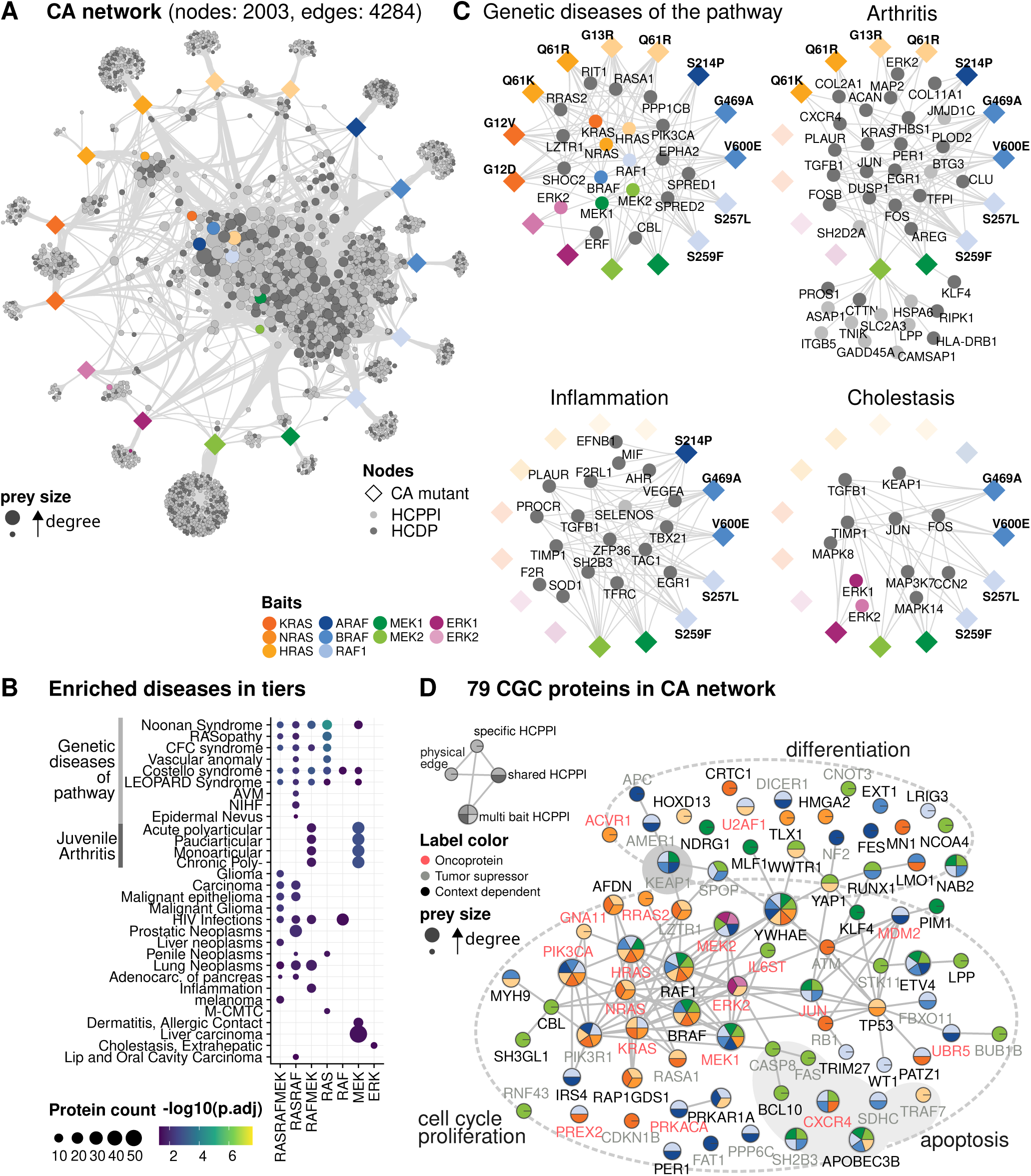
The CA pathway and disease. (A) Network representation of CA interactomes. HCDPs (DISGENET score > 0.66) are in dark grey; HCPPIs representing endogenous pathway proteins are color-coded. HCPPI size is proportional to node degree. Edges with similar start and end nodes were bundled in Cytoscape using default parameters to emphasize network topology. (B) DISGENET enrichment analysis of combined CA AP-MS and TbID-MS interactomes. HCPPIs aggregated at the tier level and shared by more than one tier are shown. Dot size is proportional to the number of proteins in each DISGENET term; color scale = - log10(adj. p-value). (C) Network representation of HCPPIs involved in selected DISGENET terms. Baits with more than three edges are highlighted for clarity. (D) STRING physical network (minimum score 0.4). CA HCPPIs annotated as oncoproteins (red) or tumor suppressors (grey) or context-dependent (black) in the CGC database. Visualized as in Figure 4B.

Interactors shared by RAS/RAF/MEK, RAS/RAF and RAF/MEK harbored HCDPs enriched in genetic anomalies of the pathway, cancers, and in HIV infection. RAF/MEK HCPPI were prominently associated with inflammatory conditions, e.g. RAF HCDPs with HIV infection, and MEK HCDPs with arthritis, dermatitis and liver carcinoma. ERK HCDPs were enriched only in extrahepatic cholestasis (Figure 5B). Zooming into paralog- and mutations-specific disease association, we observed RAS paralog-specific disease associations (e.g. KRAS G12V with endometriosis) and fine-mapped the association of HIV infections to RAF1 S257L, of inflammation, myocardial ischemia and cholestasis to MEK1DD, and of cholestasis to ERK1 (Figure S5D). Importantly, the interaction between the CA pathway and HCDPs was not limited to the paralogs showing significant enrichment in overrepresentation analysis (ORA). Rather, complex interactions emerged between multiple pathway components and HCDPs associated with specific diseases, supporting the notion that pathway activation plays a role in these conditions (Figure 5C).

The broad range of neoplasms-associated terms in the ORA (Figure 5B) suggested the existence of a core network of cancer-associated HCDPs in the pathway interactome. Intersecting the CA interactomes with the Sanger cancer gene census (CGC) revealed a network comprising 79 HCPPI and 180 interactions, 65% previously undescribed. The network featured oncogenes and tumor suppressors regulating cell cycle, apoptosis, and differentiation (Figure 5D). Thus, members of the pathway engage in concerted interactions with HCDPs involved in genetic and acquired diseases, and with a core subset of cancer-associated proteins common to a wide spectrum of neoplasms.

### Simultaneous detection of paralog and mutation-specific interactors reveals unique impact of NRASQ61R on AKT activation

RAS interactomes broadly recapitulated the canon of RAS effectors, including the expected LOI or GOI caused by CI and CA mutations.^31^ Of all the CA-RAS mutants, NRASQ61R showed the clearest preference for PI3K subunits in both the AP-MS and TbID-MS sets. This was specific, as the interaction with RAF1 and BRAF was similar across CA mutants (Figure S6). NRASQ61R interactomes also contained the scaffold IRS2, which links RTK activation to PI3K, and PI3K downstream targets including AKT3 and DEPTOR (Figure 6A). This pattern was confirmed by V5 and Streptavidin pull-downs. Additionally, the expression of DEPTOR, which modulates PI3K signaling by acting on mTOR-containing complexes both upstream and downstream of AKT,^62^ was increased in NRASQ61R lysates (Figure 6B).

**Figure 6.**
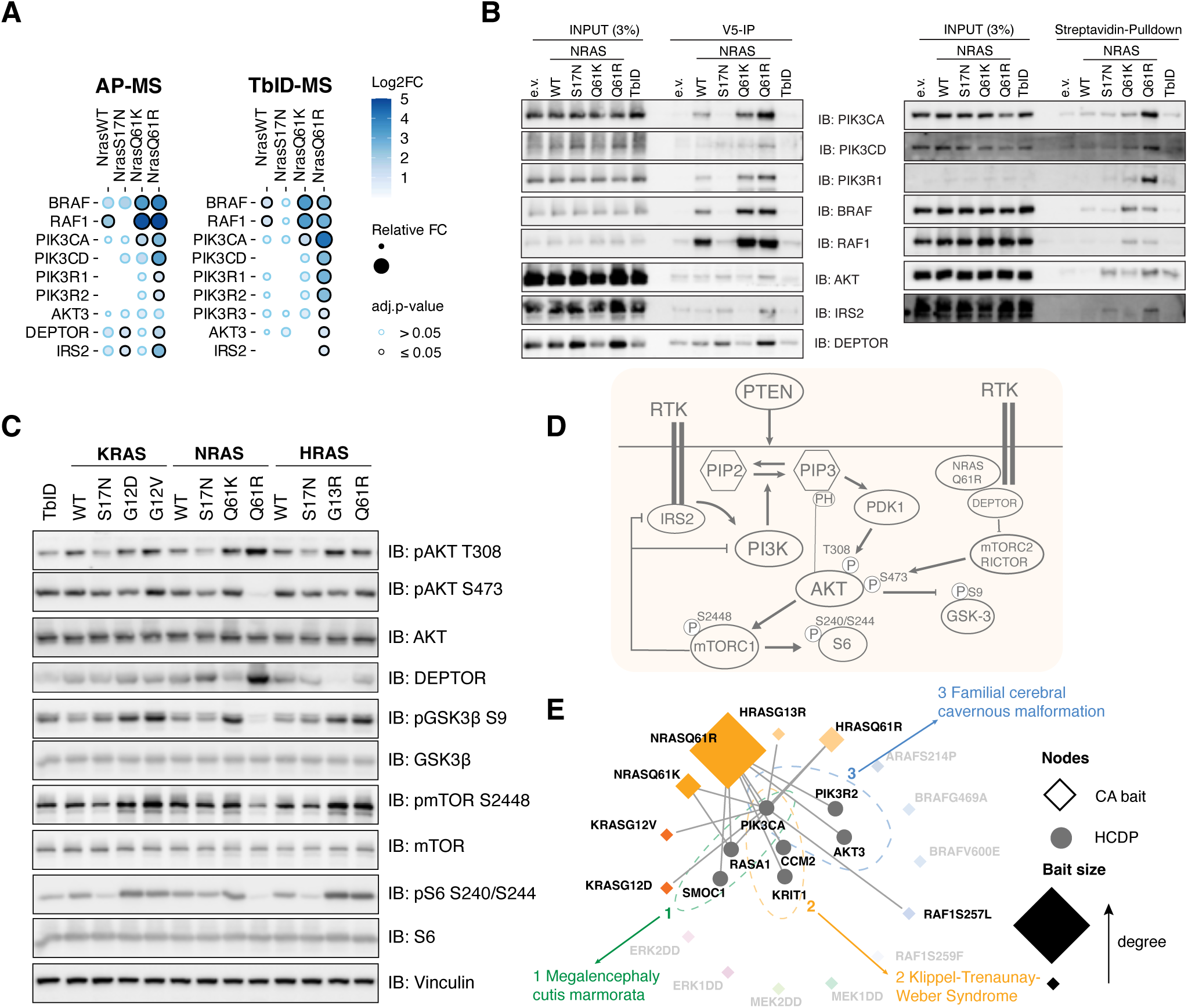
Paralog and mutation-specific effect of NRASQ61R on AKT signaling. (A) RAF and PI3K proteins interacting with NRAS. Fold change (log2 or relative) compared to control and adj. p values are shown. (B) Immunoblot validation of HCPPIs shown in A. (C) Immunoblot analysis of AKT signaling. Lysates of cells expressing all RAS paralogs in CI, WT and CA states were probed with the phosphospecific antibodies indicated on the right. (D) Schematic representation of the model described in the text. RTK = receptor tyrosine kinase. E) Network of PROS-related CA-RAS HCPPIs. Differential edges (log2FC > 1 and adj.p-value < 0.05 compared to WT) are shown. The size of the CA nodes is proportional to node degree.

NRASQ61R cell lines exhibited a distinctive AKT phosphorylation signature—high pT308 (PDK1/PI3K-dependent activation site) paired with low pS473 (mTORC2-dependent site required for full-fledged AKT activation^63^). This pattern was not observed in NRASQ61K or HRASQ61R lines (Figure 6C–D). These data support a model in which increased recruitment of the mTOR inhibitor DEPTOR by NRASQ61R would dampen mTORC2 activity, thereby reducing AKT pS473. The consequent decrease in AKT substrate phosphorylation (e.g., GSK3B and mTOR) would in turn lower mTORC1 output, read out as reduced S6 phosphorylation. Low mTORC1 activity would also alleviate the negative feedback imposed by mTORC1 on IRS and PI3K, elevating PI3K signaling and AKT pT308, consistent with our observations. Additionally, since mTOR and S6K1 phosphorylate DEPTOR and prime it for degradation, low mTOR/S6K1 activity would also stabilize DEPTOR, potentially explaining its increased abundance in our system.^62^

Thus, comparing multiple CA paralog interactomes in the same experiment allowed us to draw conclusions that could not have been derived by the comparison of individual experiments carried out in different labs and under different conditions.

In terms of disease, NRASQ61R was also the CA-RAS interacting with most proteins linked to three benign, but incapacitating *PIK3CA*-related overgrowth spectrum development diseases^64^ (PROS; Figure 6E). Elevated AKTpT308 has also been found in a mouse model of highly malignant multiple myeloma driven by germinal center B-cells-restricted *NRAS Q6*1R and *MYC* expression.^65^ Intriguingly, overexpression of DEPTOR, mostly regarded as a tumor suppressor, is necessary for cell survival in multiple myeloma,^66^ suggesting a potential link to cancer biology and the design of specific therapies.

### Crosstalk with mRNA metabolism and WNT signaling

ZFP36/TTP is a crucial regulator of the stability of AU-rich elements (ARE)-containing mRNAs, most of which code for cytokines involved in inflammation. TTP regulation, including via phosphorylation by MAPK including ERK,^67,68^ is essential to coordinate cytokine expression during inflammation.^69^ TTP can also act on its own mRNA, engendering a feedback loop important in the resolution of inflammatory processes.^68,69^ Additionally, TTP has been implicated in tumor suppression, by regulating both oncogenic and tumor suppressor transcripts.^70^

TTP was a prominent HCPPI of CA-BRAF, RAF1, MEK1 and MEK2 (Figure 7A), validated by V5 and Streptavidin pulldowns (Figure S7A). Expression of a GFP reporter containing the TNF ARE destabilizing sequence, as well as TTP itself, were strongly increased in CA-RAF and CA-MEK cell lines, indicating reduced TTP activity. This phenotype was rescued by inhibitors (Figure 7B). Importantly, RT-qPCR assays measuring endogenous TTP targets relevant in inflammation, tumor suppression, and TTP itself, yielded similar results (Figure 7C). Thus, CA-RAF and CA-MEK control mRNA turnover by regulating TTP expression and activity. Suppressing the latter, possibly via ERK-mediated phosphorylation, ^67^ resulted in the accumulation of TTP target mRNAs.

**Figure 7.**
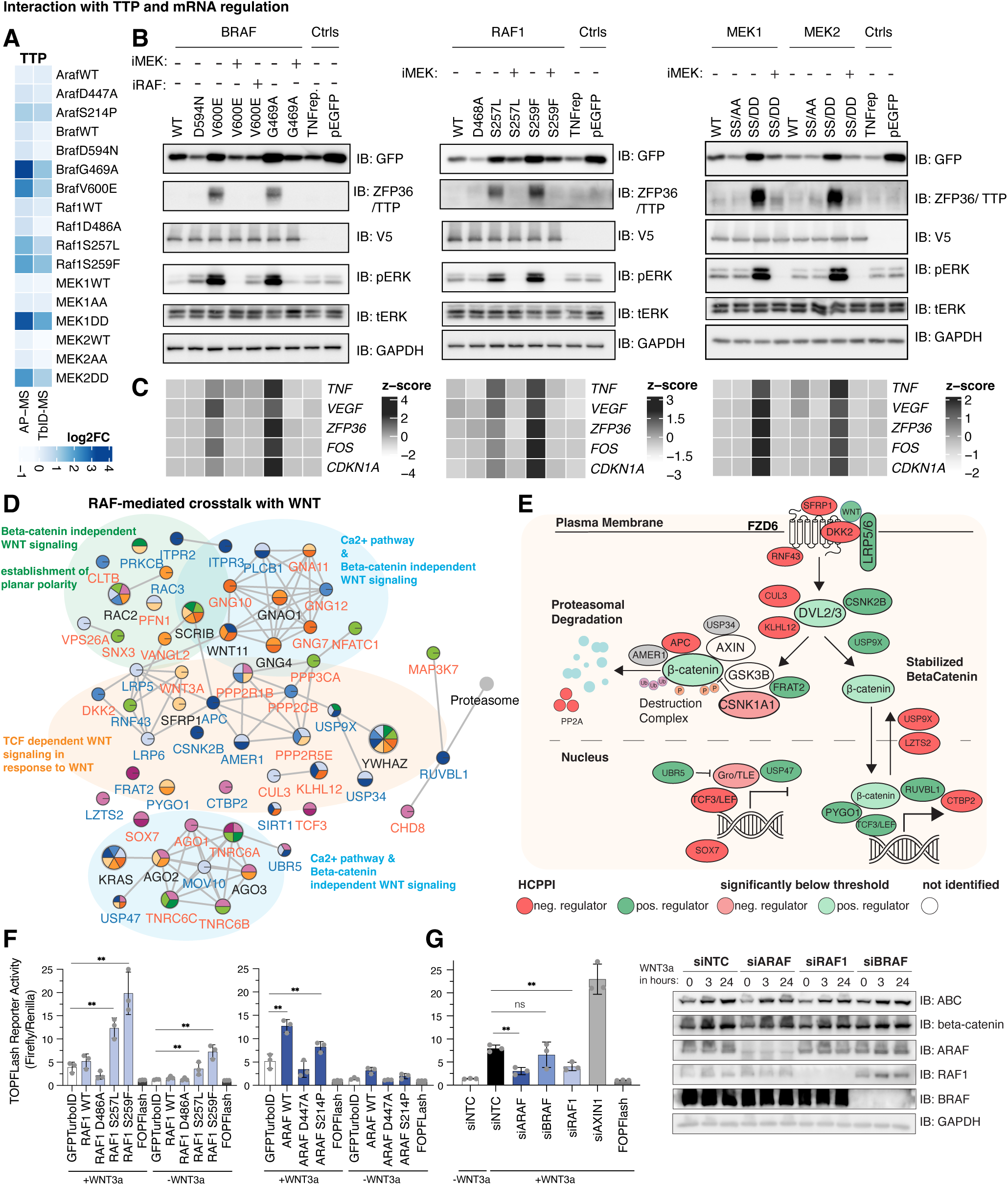
Cross-talk with mRNA metabolism and WNT. (A) Relative quantification of the abundance of TTP captured by RAF and MEK in AP-MS and TbID-MS. (B) Immunoblot analysis of RAF and MEK cell lines transfected with the GFP-TNF-ARE reporter. iRAF = dabrafenib (2.5µM, overnight); iMEK = trametinib (1µM, overnight). (C) qRT-PCR analysis of TTP targets in RAF and MEK cell lines treated as in (B). (D) STRING network of WNT-related HCPPIs. Pie size is proportional to the number of interactions with the baits (node degree). Blue labels = nodes detected by AP-MS; red = detected by TbID; black = detected by both. Edges represent physical interactions (minimum score 0.4). (E) Overview of WNT signaling featuring HCPPIs from this screen colored according to their effect on WNT/TCF signaling: green=positive; red = negative; grey = context-dependent. Core elements of the WNT pathway detected below threshold are shown in pale green or red, the missing ones are in white. (F, G) WNT/TCF-driven transcription in RAF-expressing expressing (F) or RAF-depleted (G, left panel) cell lines determined by TOPflash luciferase assay. Cells were left untreated or treated with WNT3A (100 ng/ml, overnight). Results are plotted as mean ± SD. \**p ≤* 0.05; \*\**p ≤* 0.002, according to Student’s *t* test. (G, right panel) Immunoblot analysis Active β-CATENIN (ABC) levels in RAF-proficient and deficient cells, treated or not with WNT3a for the indicated times.

We next queried Reactome for signaling pathways enriched in our datasets. Excluding general entries (e.g. “signaling by RHO GTPases”, “RTK pathways”), “β-CATENIN-independent WNT signaling” ranked highest, surpassing pathway-specific entries like “signaling by BRAF mutants” or “Signaling to ERK”. The entries “Signaling to WNT” and “TCF-dependent signaling in response to WNT” ranked lower but were still significantly enriched, at levels comparable to “FGF signaling”. Pathway interactors present in “<u>Signaling</u> <u>by WNT</u>” organized into two hubs—WNT3A, linked to canonical TCF-dependent signaling, and WNT11, associated with non-canonical β-catenin-independent signaling^71^ (Table S2; Figure 7D-E). The latter contained proteins involved in the WNT/Ca2^+^ pathway and in the establishment of planar cell polarity during development. We also observed a RISC cluster, involved in the translational control of NLK, a kinase repressing the expression of β-CATENIN targets. The WNT3A cluster contained mainly proteins that modulate TCF-dependent signaling at multiple levels (Table S2). Positive regulators included the co-receptors LPR6 and 5, and proteins promoting β-CATENIN- and TCF-induced transcription; several HCPPIs had complex roles in WNT signaling, e.g. modulating the stability of β-CATENIN, DVL or AXIN. The interactomes also included proteins antagonizing WNT binding to its receptor or receptor/co-receptor interaction, and inhibitors of β-CATENIN- induced gene expression. A strong cluster represented by the proteasome was involved in the control of both β-CATENIN-independent und TCF-dependent signaling.

Consistent with the patterns observed in the subcellular networks, state-specific analysis showed that the CA network had the strongest connectivity and the highest node count. De novo additions were predominantly peripheral receptors/adaptors and extra signaling paralogs, typically contributed by single baits. The fraction of nodes supported by multiple baits increased (Figure S7C).

Thus, ERK pathway components interacted both with positive regulators of β-CATENIN-independent WNT signaling and with context-dependent or negative regulators of TCF-dependent signaling. While some WNT core elements were either not detected (AXIN), or were below threshold (FZD6, DVL2/3, β-CATENIN) in our interactomes, the co-receptors LRP5/6 were HCPPI of WT and CA-RAF1. This interaction was validated by V5 pulldowns, which also showed less LRP5/6 in CI-RAF1 (Figure S7B). CA-RAF1 promoted expression of a TCF/LEF reporter even in untreated cells, and strongly increased expression in WNT3A-treated cells. The effect of ARAF, which was connected to AMER1 and APC, was most evident in WT-ARAF, WNT3A-treated cells (Figure 7F). Depletion of RAF paralogs using specific siRNAs confirmed that ARAF and RAF1, but not BRAF, contribute to WNT-induced reporter expression and to the accumulation of active β-CATENIN (Figure 7G), suggesting an involvement of ARAF and RAF1 in the initial steps of WNT signaling.

## Discussion

### A comprehensive pathway-centric approach generates a rich resource for biomedical research

We combined AP-MS, which captures proteins in stable complexes at the time of isolation, with TbID-MS, which detects transient/vicinal interactors, to comprehensively profile the interactomes of all ERK pathway components in WT and perturbed states. This strategy recovered the largest set of high-confidence PPIs (HCPPI) for the ERK pathway reported to date (Figure 1), creating a resource with great discovery potential. AP-MS and TbID-MS provided complementary, orthogonal evidence that validated and expanded the network, while the quantitative information generated by DIA-MS defined state-agnostic and state-specific HCCPI. The pathway interacted vigorously with components of other signaling and metabolic pathways and with compartment-specific proteins. Activation increased both the number of HCPPIs per bait and the convergence of the pathway’s interactomes.

Interrogation of the system-level RAStoERK atlas delivered state-resolved, pathway-wide networks that share a common topology, with consolidated cores comprising nodes contributed by more than one prey and dynamic, context- and bait-specific peripheral nodes typically gained in the CA networks. These networks support the conclusion that signaling crosstalk and spatiotemporal coordination are pathway-level phenomena. They distinguish core dependencies from peripheral vulnerabilities, and provide a basis for testable mechanistic hypotheses, systematic dissection of pathway crosstalk, and informed prioritization of intervention points.

The HCPPIs of CA mutants also comprised many HCDPs, converging mostly at the intersection of the RAF/MEK tiers. HCDPs were involved in clinical conditions including genetic diseases of pathway components, but also inflammation-related diseases; and in the context of cancer, the combined datasets revealed a core network of oncogenes and tumor suppressors whose nodes were contributed by more than one bait.

None of these conclusions could have been drawn by compiling interactomes generated in different labs and under different conditions.

The experimental framework we used has at least three intrinsic limitations. HEK293 cells do not fully recapitulate in vivo physiology, the baits are expressed alongside endogenous counterparts, and AP-MS/TbID-MS each have method-specific biases. We chose HEK293 cells for robustness, ease of handling, and alignment with large public interactome resources generated in the same system. ^12,13,14,55^ Rather than endogenously tagging the bait loci, we chose to maximize comparability by generating inducible polyclonal cell lines and titrating our baits’ expression to the level of the corresponding endogenous protein. This avoided the uncontrollable, undetectable clonal variation generated by the multiple rounds of single cell cloning required for endogenous targeting of both bait alleles plus the introduction of point mutations, which would have jeopardized comparisons across 35 cell lines. Finally, parallel deployment of AP-MS and TbID-MS, which capture physical complexes and proximal neighbors respectively, should balance method-specific sensitivities and biases, such as over- and underrepresentation of proteins/peptides based on their chemical properties or, in the case of TbID-MS, protein size and amino acid composition. This dual-technology, quantitatively anchored design mitigates these constraints and enables reproducible, pathway-wide discovery. Our one-lab/one-pathway proteomic approach has uncovered paralog- and even mutation specific interactors, and previously undescribed interactors that opened avenues into crosstalk with unrelated signaling cascades impacting inflammation or development. These results validate our approach and highlight the value of the comprehensive, pathway-centric datasets available at *RAStoERK.univie.ac.at* for both hypothesis-free and hypothesis-driven interrogation of the pathway, with the potential to uncover therapeutically actionable mechanisms. Future research will focus on identifying how cancer-associated mutants, chemical RAF and MEK inhibitors, or degraders rewire the network, and integrating this information with changes in the proteome/phosphoproteome. The results could pave the way for the discovery of actionable vulnerabilities in the pathway rewired by oncogenes or inhibitors.

## Acknowledgments

We thank the Mass Spectrometry Facility, specifically N. Hartl and W. Chen, who performed the proteomics analysis with assistance of the VBCF instrument pool. The skillful technical assistance of Karin Ehrenreiter is greatly appreciated. We thank the Menche Lab, Dr. Thomas Decker, Dr. Gijs Versteeg (both Max Perutz Labs) and Dr. Ulrich Stelzl (University of Graz) for insightful discussions. This work was supported by FWF grant P35464 (to M.B.).

## Author contributions

Conceptualization, G.V. and M.B.; methodology, S.D.; formal analysis, S.D.; investigation, G.V., M.H., L.C, A.S.; writing review and editing, M.B., G.V., S.D., J.M.; visualization, M.B., G.V., S.D.; supervision, M.B. and J.M.

## Conflict of interest

The authors declare no competing interests.

## Materials availability

All unique/stable reagents generated in this study are available from the lead contact with a completed materials transfer agreement.

## Data and code availability

- Raw MS data have been deposited to the ProteomeXchange Consortium via the PRIDEpartner repository. All data are publicly available; see accession numbers in the key resource table.
- All original code has been made available in the supplement (Data S1) and will be deposited to GitHub and Zenodo and will be publicly available as of the date of publication.
- The webplatform is available at <u>RAStoERK.univie.ac.at</u> and will be publicly available as of the date of publication.

Any additional information required to reanalyze the data reported in this work is available from the lead contact upon request.

## STAR Methods – Materials and Methods

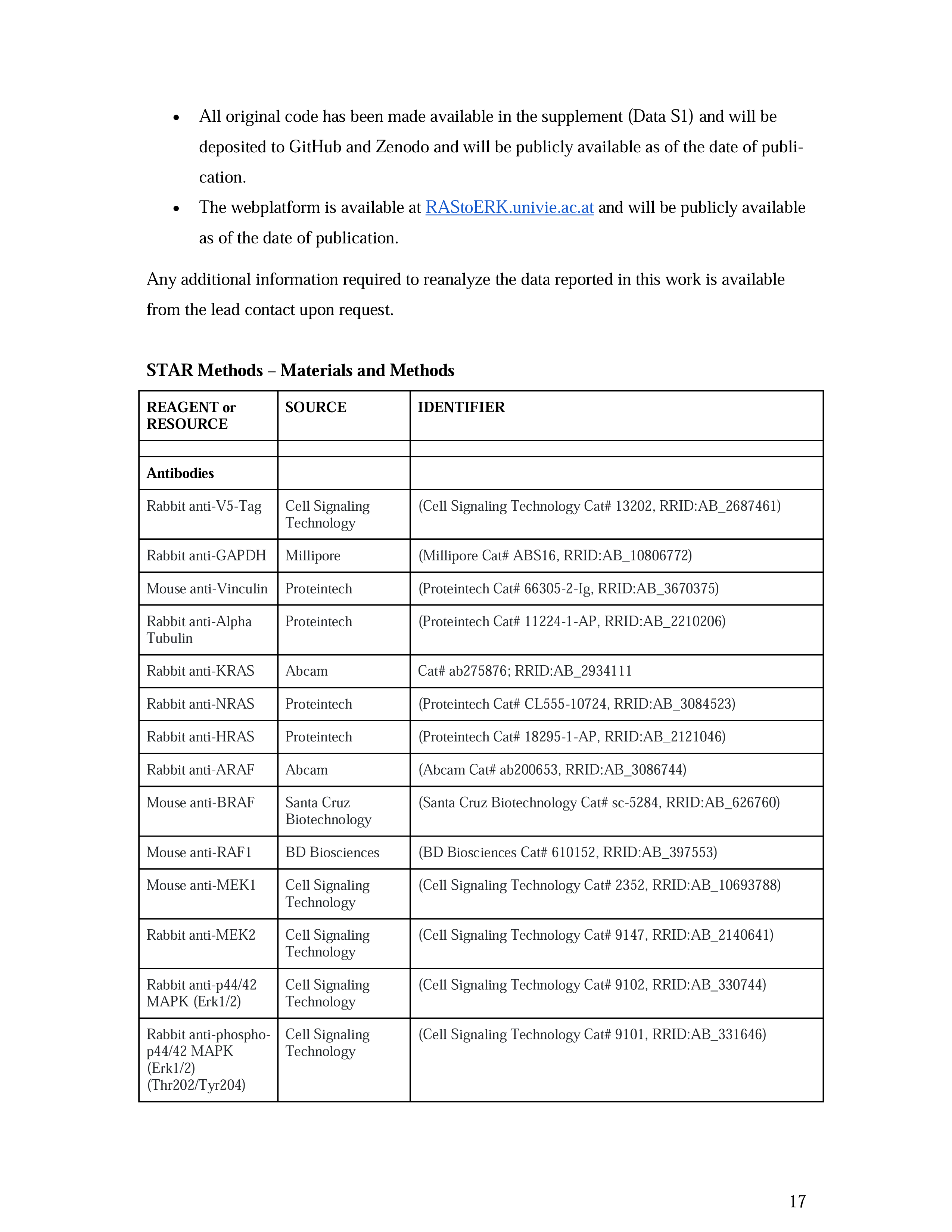
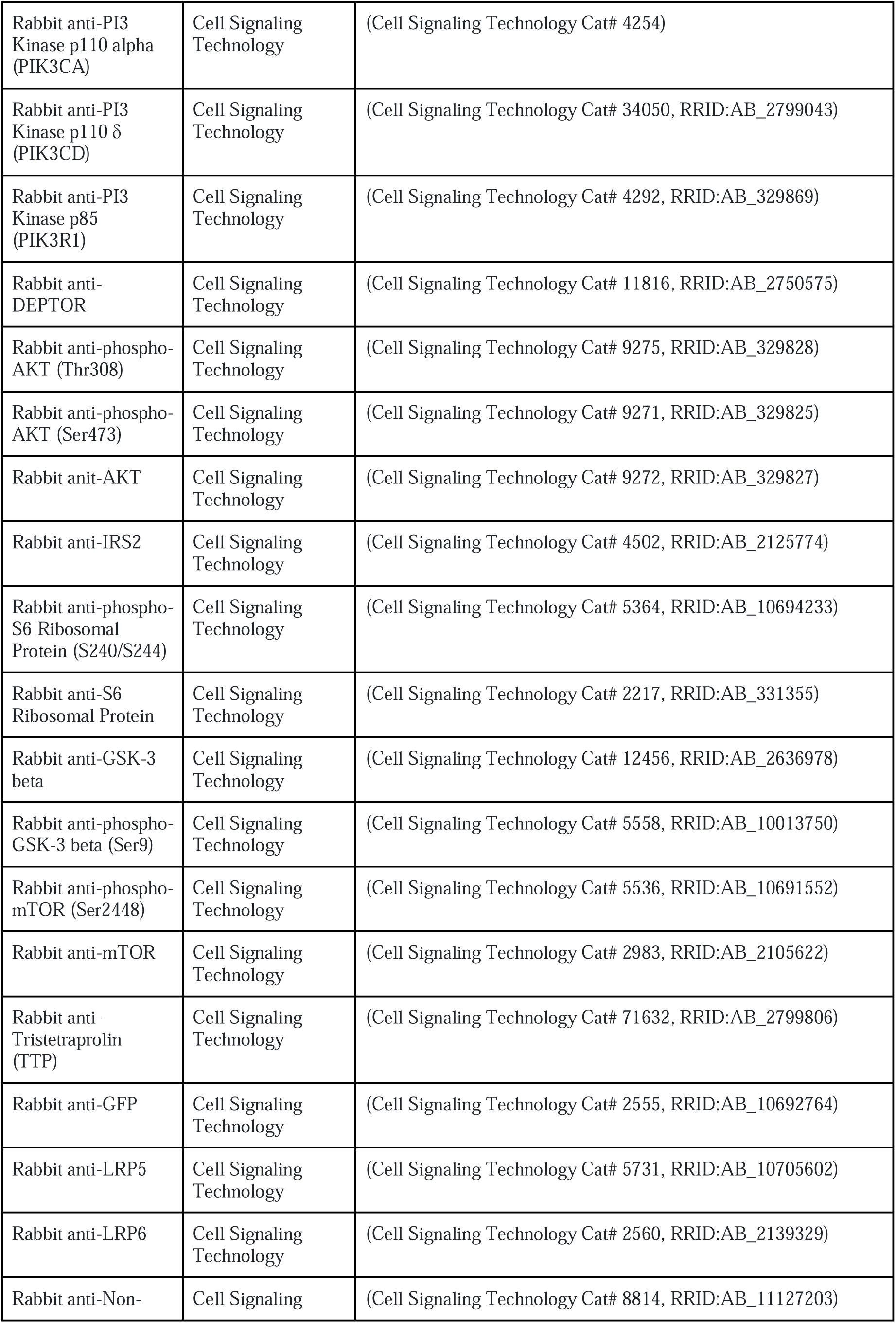
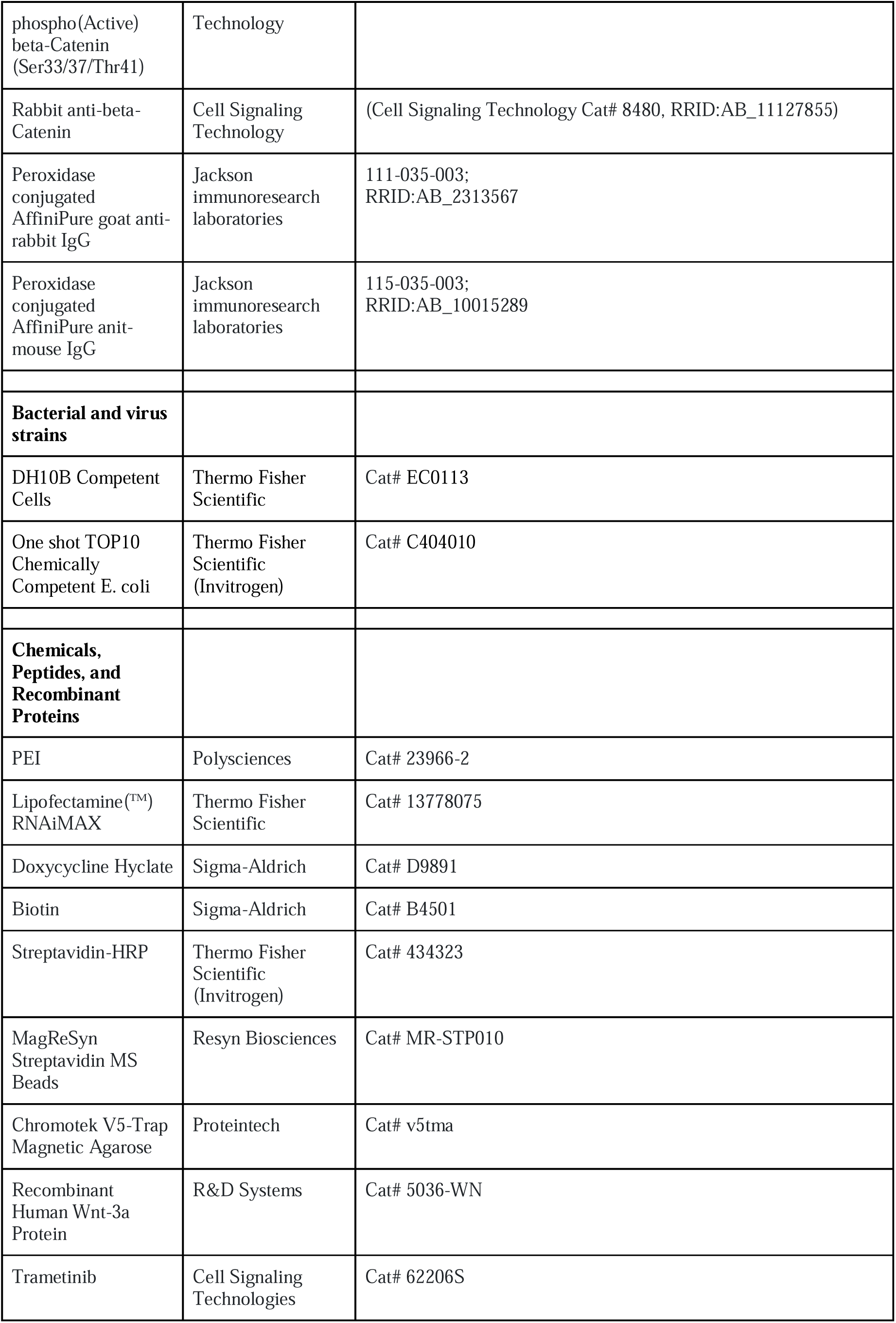
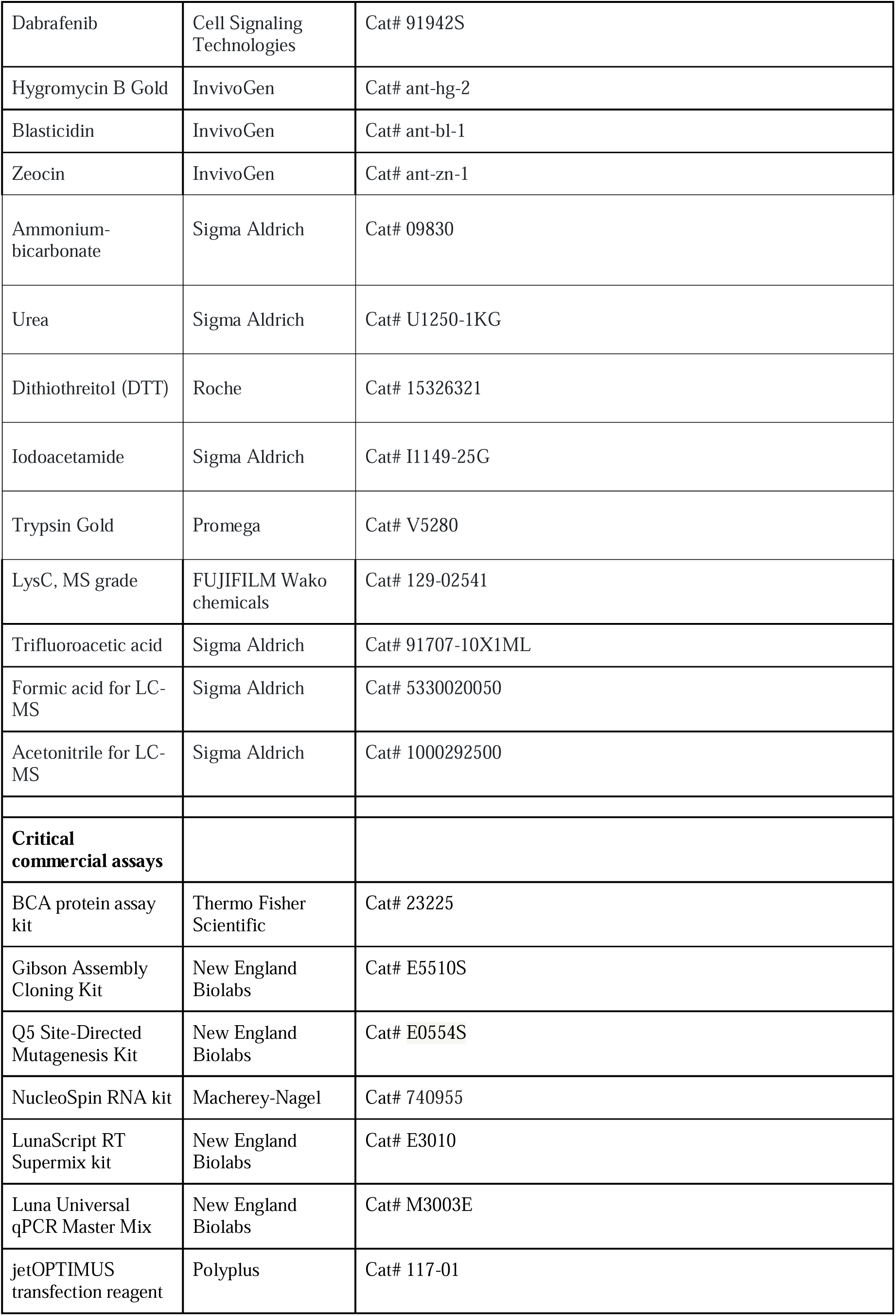
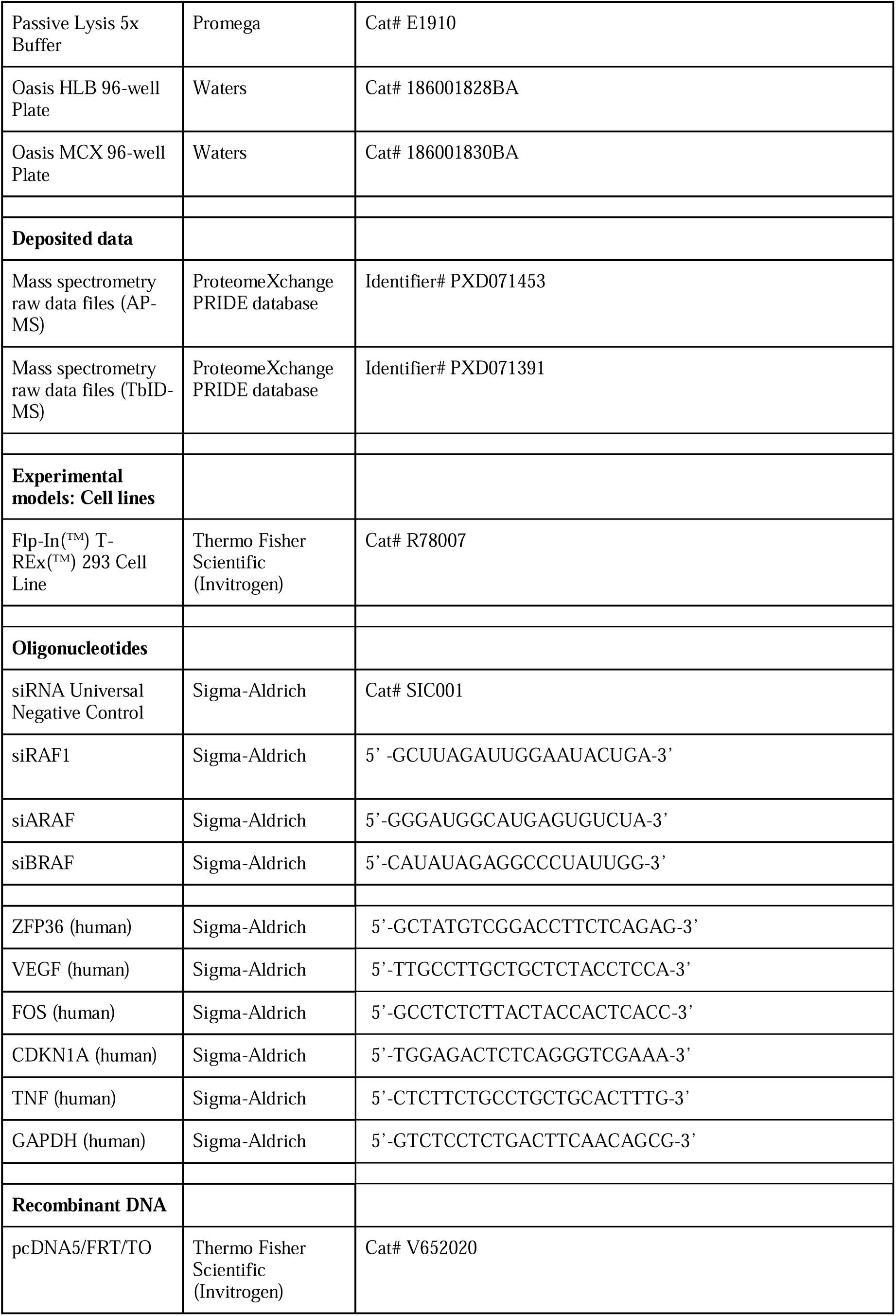
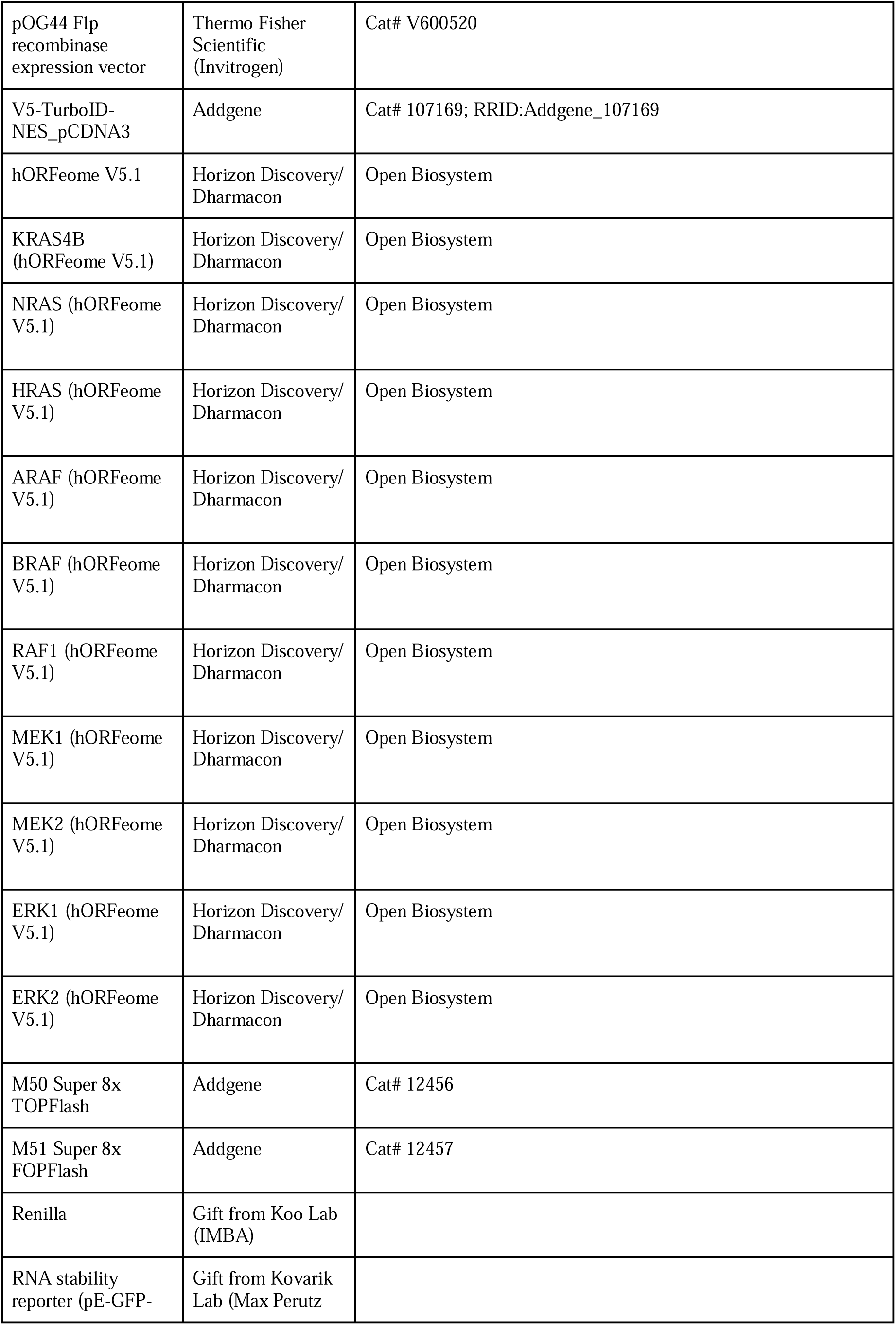
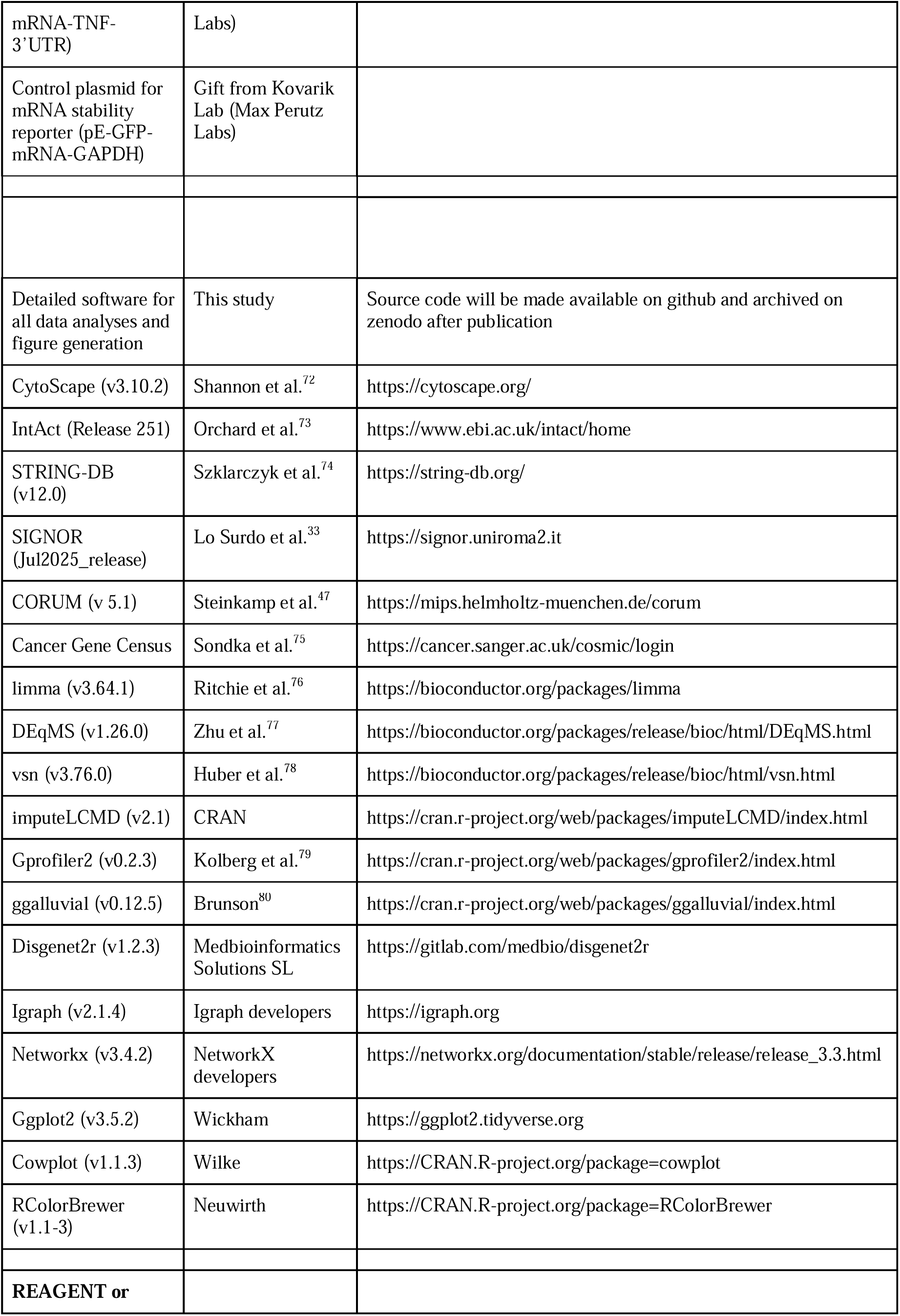
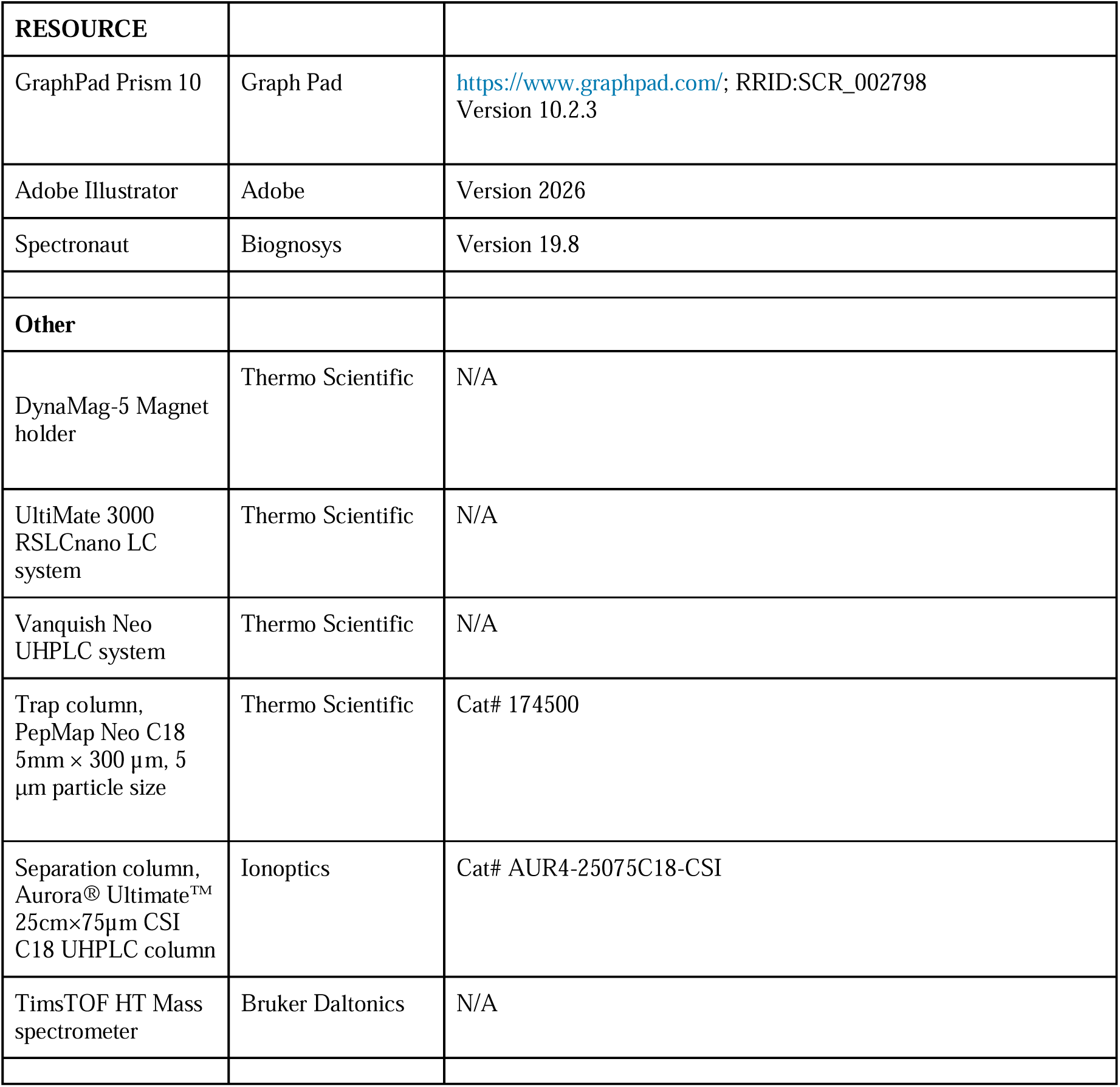

### EXPERIMENTAL MODEL AND STUDY PARTICIPANT DETAILS

#### Cell culture

Flp-In T-REx HEK293 cell lines (#R78007, Invitrogen) were cultured in Dulbecco’s modified Eagle’s medium (Sigma-Aldrich, D6429), supplemented with 10% fetal bovine serum (Sigma), 1% penicillin/streptomycin at 37°C in a humidified incubator with 5% CO_2_. Cells were tested negative routinely for mycoplasma contamination by PCR.

### METHOD DETAILS

#### Cloning and cell line generation

Complementary DNAs (cDNA) of each pathway member were obtained from the human ORFeome collection (hORFeome v5.1). For the generation of expression constructs, cDNAs of the pathway components were cloned using the Gibson Assembly Cloning Kit (NEB, E5510S) into a doxycycline-inducible N- or C-terminal V5-TurboID tagged vector pcDNA5™/FRT/TO (Thermo Fisher Cat. No. V601020) for mammalian expression.

Point mutations were then introduced into the WT pathway components via site-directed mutagenesis using the Q5 Site-Directed Mutagenesis Kit Protocol (NEB, E0554), according to the manufacturers protocol. The most common cancer-associated somatic mutations were chosen from https://www.cbioportal.org querying curated sets of non-redundant studies; and from the cancer hotspots site https://www.cancerhotspots.org. For KRAS, we focused on KRAS4b, which represents about 90% of the KRAS proteins in cancer, and on its cancer hotspot G12D; and on the Q61R mutation of NRAS and HRAS.

Ligated plasmid product were transformed into competent cells (NEB 5alpha competent *E.Coli* Cat.No.: C2987l, *E. coli* One Shot TOP10 chemically competent Thermo Fisher Scientific Cat. No.: C404010)

All plasmids were verified by restriction digest analysis, followed by insert sequencing, as well as full plasmid sequencing (Microsynth).

#### Generation of stable cell lines

For the generation of stable isogenic cell lines, Flp-In T-REx HEK293 cells were co-transfected with the vector pOG44 (Invitrogen), encoding the Flp-recombinase and the pcDNA5™ 5/FRT/TO plasmids that harbor the incorporated wild-type, inactive or active mutants using jetOPTIMUS. 72 hours post transfection, cells were transferred into a new vessel containing selection media (15µg/mL blasticidin and 100 µg/ml hygromycin). Selection media was replaced and cells were split when necessary until non-transfected cell-lines were dead (roughly 7-14 days). The surviving cells were expanded, frozen. The expression of the bait proteins was tested and validated via western blot analysis.

#### RT-qPCR

Culture dishes containing the cells were removed from the incubator and immediately placed on ice, followed by two washes with ice-cold PBS. The total mRNA was isolated with the nucleospin RNA kit (Macherey-Nagel, Cat# 740955) according to the manufacturer’s instructions. RNA concentrations were measured using NanoDrop(™) 2000 Spectrophotometer and 1µg of RNA per sample was subsequently used for the cDNA synthesis reaction. cDNA was synthesized using the LunaScript RT Supermix kit (NEB Cat# E3010). The samples were diluted 1:10 with ddH2O (nuclease free). RT-qPCR was performed on a BioRad CFX96 touch real-time PCR detection system according to the manufacturer’s protocol (Luna Universal qPCR Master Mix, M3003E, NEB).

#### Transfection

Transfections for DNA were carried out using jetOPTIMUS (Polyplus), or PEI following manufacturer’s instructions. For transfection of siRNAs, Lipofectamine RNAiMAX (Invitrogen) for siRNA according to manufacturer’s instructions. siRNAs were used at 30 nM final concentration and cells were lysed 48h post transfection.

#### Reporter Assay

M50 Super 8x TOPFlash or M51 Super 8x FOPFlash (as negative control) plasmids were cotransfected with Renilla luciferase (internal control for transfection efficiency) using PEI (1:2 DNA to PEI ratio). Cells were stimulated with recombinant Wnt3a (100ng/mL) overnight. Cells were lysed in 1x Passive Lysis Buffer (Promega E1910).

#### Cell lysis, immunoprecipitation and western blotting

Cells were lysed in ice-cold 10 mM Tris-HCl (pH 7.4), 137 mM NaCl, 1mM CaCl2, 1mM MgCl2, 1% NP-40, 1 mM Na3VO4, 50 mM NaF, 2mM PMSF, and protein inhibitor cocktail (Roche). The lysates were placed on a rotation wheel at 4 °C for 10 min. Clearing of debris was performed by centrifugation at 15,000 x g for 10min at 4 °C. Protein concentrations were determined using Pierce® BCA Protein Assay Kit according to the manufacturer’s protocol and measured on a Synergy H1 microplate reader. Lysates were adjusted to contain equal concentration of protein and used for immunoprecipitation (V5 or Streptavidin pulldown) or Western blot analysis.

#### Immunoblotting

Immunoprecipitates or cell lysates were subjected to 8-15% SDS-PAGE followed by overnight transfer (Biorad Trans-Blot Cell) for 18h (200 mA for 16h, 400 mA for 2h) at 4°C onto a 0.45µM PVDF membrane (Immobilon-P; Millipore). The blots were blocked (5% milk in TBST for 30-60min) and probed for 1h or o/n at 4°C with indicated primary antibody, followed by incubation with secondary antibody for 1h at RT. Protein detection was performed using chemiluminescent substrate (SuperSignal West Pico PLUS, Thermo Fisher Scientific, Cat# PI34580). Images were acquired using Chemidoc touch gel imaging system (Biorad). For stripping, membranes were washed with ddH_2_O for 5 minutes, subsequently cooked for 15 minutes in a microwave at 800W upside down in ddH_2_O.

#### Affinity Purification sample preparation for mass spectrometry (AP-MS) or Western blot

Experiments were performed in biological triplicates, harvested from 10cm cell culture dishes with a confluency of 80-90%. All cell lines were placed in culture at the same time and seeded for the experiment. Once the cell lines reached 60-70% confluency, doxycycline was added to the media for bait-protein expression overnight. For affinity purification, one confluent 100 mm tissue culture plate was harvested per replicate. Cells were washed 3x with ice-cold PBS and flash frozen in liquid nitrogen. The cell pellets were lysed in a buffer containing 20 mM Tris-HCl (pH 7.4), 137 mM NaCl, 1 mM CaCl2, 1 mM MgCl2, 1% NP-40, 1 mM Na3VO4, 50 mM NaF, 2 mM PMSF, and protein inhibitor cocktail (Roche). The cleared cell lysate was incubated with ChromoTek V5-Trap® Magnetic Agarose (Proteintech #V5tma) for 1 h on an orbital shaker at 4 °C. Following incubation, the affinity matrix was washed 3x with lysis buffer and 6 times with PBS.

#### TurboID sample preparation for mass spectrometry (TurboID-MS) or Western blot

24 hours prior to the harvesting of cell lines at 80% confluency, doxycycline was added to the media for bait-protein expression. 10 minutes before lysis, biotin (Sigma Aldrich Cat. No.: B4501) was added to a final concentration of 500 µM. One confluent 100 mm tissue culture plate was harvested per replicate. Cells were washed 3x with ice-cold PBS and flash frozen in liquid nitrogen. The cell pellets were lysed in RIPA Buffer containing 50 mM Tris-HCl, pH7.4,, 150 mM NaCl, 1% NP-40, 0.1% SDS, 0.5% sodium deoxycholate supplemented with 1 mM Na3VO4, 50 mM NaF, 2 mM PMSF, and protein inhibitor cocktail (Roche). The cleared cell lysate was incubated with MagReSyn® Streptavidin MS Beads for the high capacity capture of biotinylated proteins (MR-STP010 MS Wil) for 1 h on an orbital shaker at 4 °C. Following incubation, the affinity matrix was washed 3x with lysis buffer and 6x with ice-cold PBS.

#### Mass Spectrometry

##### Sample preparation for mass spectrometry analysis

###### APMS

The beads were transferred to a new tube and resuspended in 30 µL 2 M urea and 50 mM ammonium bicarbonate and digested with 75 ng LysC (mass spectrometry grade, FUJIFILM Wako chemicals) and 75 ng trypsin (Trypsin Gold, Promega) at room temperature for 90 min. The supernatant without beads was transferred to a new tube. The beads were washed with 30 µL 1 M urea and 50 mM ammonium bicarbonate, and the supernatants were. Disulfide bonds were reduced with 2.4 µL of 250 mM dithiothreitol (DTT) for 30 min at room temperature, followed by the addition of 2.4 µL of 500 mM iodoacetamide and incubation for 30 min at room temperature in the dark. The remaining iodoacetamide was quenched with 1.2 µL of 250 mM DTT for 10 min. 50 mM ammonium bicarbonate was used to dilute the urea concentration to 1M. The proteins were digested with 75 ng LysC and 75 ng trypsin at 37°C overnight. The digest was stopped by the addition of trifluoroacetic acid (TFA) to a final concentration of 0.5 %, and the peptides were desalted using an HLB 96-well plate (Waters), followed by desalting with an MCX 96-well plate (Waters), following the manufacturer’s protocol.

###### TurboID

The beads were transferred to a new tube and resuspended in 50 µL 1 M urea and 50 mM ammonium bicarbonate. Then, disulfide bonds were reduced with 2 µL of 250 mM dithiothreitol (DTT) for 30 min at room temperature, followed by the addition of 2 µL of 500 mM iodoacetamide and incubation for 30 min at room temperature in the dark. The remaining iodoacetamide was quenched with 1 µL of 250 mM DTT for 10 min. The proteins were digested with 300 ng trypsin (Trypsin Gold, Promega) at 37°C overnight. The digest was stopped by the addition of trifluoroacetic acid (TFA) to a final concentration of 0.5 %, and the peptides were desalted using an HLB 96-well plate (Waters), followed by desalting with an MCX 96-well plate (Waters), following the manufacturer’s protocol.

##### Liquid chromatography-mass spectrometry analysis

Liquid chromatography for LC-MS analysis was performed on an UltiMate 3000 RSLCnano LC system (Thermo Scientific) or a Vanquish Neo UHPLC system (Thermo Scientific), respectively. Both LC setups were coupled to a timsTOF HT (Bruker), equipped with a CaptiveSpray ion source (Bruker), and a Butterfly Portfolio Heater (Phoenix S&T). About 500ng peptides were loaded onto a trap column (PepMap Neo C18 5mm × 300 µm, 5 μm particle size, Thermo Scientific) using 0.1% TFA as mobile phase, and separated on an analytical column (Aurora Ultimate XT C18, 25 cm × 75 µm, 1.7 µm particle size, IonOpticks), applying a linear gradient starting with a mobile phase of 98% solvent A (0.1% FA) and 2% solvent B (80% acetonitrile, 0.08% FA), increasing to 35% solvent B over 60 min at a flow rate of 300 nl/min. The analytical column was heated to 50°C.

The mass spectrometer was operated in data-independent acquisition (DIA) parallel accumulation serial fragmentation (PASEF) mode. MS2 data were acquired with eight PASEF scans per duty cycle, each containing three m/z windows. The m/z window widths were adjusted based on the expected precursor density, covering a total range of 300–1200 m/z. The ion mobility range was set to 0.64-1.42 V*s/cm, and the accumulation and ramp time were set to 100 ms. TIMS elution voltages were calibrated linearly to obtain the reduced ion mobility coefficients (1/K0) using three Agilent ESI-L Tuning Mix ions (m/z 622, 92,2 and 1,222). Collision energy for fragmentation was scaled linearly with precursor mobility (1/K0), ranging from 20 eV (at 1/K0□=□0.6□V*s/cm) to 59 eV at (at 1/K0□=□1.6□V*s/cm).

LC-MS system performance was monitored daily by analyzing a quality control (QC) standard consisting of 10 ng HeLa cell lysate digest. Key performance metrics—specifically peptide spectral matches (PSMs), median mass accuracy, and total ion chromatogram (TIC) profiles—were evaluated to ensure instrument stability. Deviation from established baselines served as criteria for instrument performance and necessary maintenance, including mass calibration.

The samples were processed and injected following a block randomization design.^81^ Each block represented one replicate for each condition, with samples in randomized order within each block. The design is documented with the raw data on the PRIDE repository.

##### Data analysis - mass spectrometry-based proteomics

#### Data availability

Mass spectrometry raw data and associated Spectronaut output tables were deposited to the ProteomeXchange Consortium via the PRIDE partner repository^82^ which will be made available after publication.

#### Proteomics data analysis

MS raw data were converted to htrms format using HTRMS converter 19.8 (Biognosys). Then it was searched with Spectronaut 19.8 (Biognosys) in directDIA+ mode against the Uniprot human reference proteome (version 2025_01, www.uniprot.org), concatenated with a database of 379 common laboratory contaminants (release 2023.03, https://github.com/maxperutzlabs-ms/perutz-ms-contaminants). The search was performed with full trypsin specificity and a maximum of two missed cleavages at a protein and peptide spectrum match false discovery rate of 1%. Carbamidomethylation of cysteine residues was set as fixed, oxidation of methionine, and N-terminal acetylation as variable modifications. The cross-run normalization was disabled, and all other settings were left at their default values.

#### Post-processing and differential abundance analysis

The Spectronaut output was analyzed using the R programming language extending the codebase from amica.^83^ Protein intensities less than 32 were set to missing to remove low quality quantification values. LFQ protein intensities were log2-transformed and normalized across samples using variance stabilization normalization (VSN) for the AP-MS dataset, and median sweeping for the TbID-MS dataset. After filtering out contaminants and proteins with fewer than two peptides or fewer than three quantified values in at least one bait group’s triplicates, the missing normalized LFQ intensity values were imputed with random values from a normal distribution using the impute.MinProb function from the imputeLCMD library. Differences between groups were statistically evaluated using DEqMS/limma. Batch effects were accounted for during statistical testing by integrating the output of limma’s duplicateCorrelation function into the linear model.^76,77^

#### Interactome analysis

All bait samples were in triplicates. Two sets of negative GFP controls were used to filter the dataset: GFP low expression (n = 9) and GFP high expression (n = 9). As a first filter, a protein needed to have a significant adjusted p-value (padj < 0.05) and a positive log2FC in the comparisons over both controls in a given bait. As a second filter, log2FC thresholds were applied using the high expression GFP controls as background. To account for different expression levels within the pathway, tier-specific log2FC thresholds were applied to the various interactomes. In general, appropriate thresholds were determined by inspecting log2FCs of known interactors from the IntAct and Signor databases in the WT interactomes.

| Tier | AP-MS log2FC threshold | TbID-MS log2FC threshold | Examples |
| --- | --- | --- | --- |
| RAS | 1 | 0.77 | PLCE1, PI3K kinases |
| RAF | 1.16 | 0.84 | CCT chaperones, PI3K kinases |
| MEK | 0.99 | 0.87 | ERK2, KSR1 |
| ERK | 0.97 | 0.84 | FGFR1, MKNK1 |
| KRAS | 0.73 | 0.77 | PI3K kinases, BRAF |

In order to ensure a high confidence interactome, an additional filtering strategy was implemented to eliminate common bystander proteins co-purifying with baits or exhibiting non-specific affinity for the affinity matrix and epitope tags.

We excluded prey proteins annotated with GO terms frequently associated with high-abundance contaminants in the CRAPome database,^84^ specifically those involved in RNA processing and translation, such as ribosomal proteins (ribosomal genesis, GO:0042254) and the spliceosome (spliceosomal complex, GO:0005681; RNA splicing, GO:0008380). Furthermore, our TbID-MS data showed non-specific enrichment of large mitochondrial complexes (mitochondrial ribosome, GO:0005761; cellular respiration, GO:0045333) and nuclear chromatin components (chromatin organization, GO:0006325). Because these proteins were consistently enriched across various plasma membrane and cytosolic baits, they were categorized as background and removed.

This stringent, data-driven filtering strategy removes not only proteins listed in the CRAPome database, but also proteins observed as consistently enriched across multiple baits in our dataset, capturing method-specific background that frequency-based CRAPome filtering would miss. Yet, it may inadvertently exclude some genuine protein-protein interactions (PPIs). Importantly, the excluded list was manually cross-referenced to ensure that no bona fide known interactors of the RAS/RAF/MEK/ERK pathway were removed.

#### Differential interactions (Mutant vs. WT)

Differential abundance analysis between the triplicates of a mutant versus the triplicates of its WT was also conducted with limma/DEqMS. Differential interactions between a CA or CI bait and its WT bait protein were only considered for the combined preys of the mutant and the WT bait. Preys having an abs(log2FC)>1 and adj. P-value < 0.05 were considered as differential.

#### Comparison with other data sets and percentage of known and novel PPIs

Individual MS data sets using the core pathway proteins as baits (the core pathway protein had the value “psi-mi:MI:0496(bait)” in the role column the IntAct table) were downloaded from the IntAct database (Release 251 - September 2025). Figure 1D focuses on the ten most extensive studies, showcasing those with the highest number of PPIs across the ten core pathway bait proteins. The percentage of known and novel PPIs was calculated using all interactions with no filters in the IntAct database.

#### Overlap coefficient

The overlap coefficient (OVL) between sets of preys for all pairwise comparisons of baits was computed using the formula OVL(A,B) = | A B |/min(|A|, |B|).

#### Kinase and phosphatase tree visualization

Kinase data was uploaded to CORAL^39^ and phosphatase data was uploaded to CORALp.^44^

#### Over-Representation and disease enrichment analysis huMAP3.0 complex analysis

The huMAP3.0 data set was downloaded and converted to a gmt file. Using g:profiler2’s ‘upload_GMT_file’ and ‘gost’ functions, enrichment analysis was conducted for each bait’s combined interactome. Given the redundancy in the huMAP3.0 resource, a graph-based approach was used to identify sub-complexes of larger protein complexes. Sub-complexes which were full subsets of larger complexes were removed from further analysis. This left only non-redundant complexes.

In the next step, a similarity graph was constructed for the remaining complexes (two complexes were connected if the overlap coefficient between the sets of their complex members was greater than 0.3). Final clusters of overlapping complexes were computed using the ‘cluster_louvain’ function from the igraph library. Finally, huMAP3.0 complexes from the same cluster were combined to a single meta-complex and visualized in Figure 3.

#### Subcellular Over-Representation Analysis

The organelle IP data set was downloaded from Hein et al., 2025 and merged with the interactome from this study. Proteins from the endomembrane organelles ‘recycling_endosome’, ‘early_endosome’, ‘lysosome’, ‘trans-Golgi’, ‘Golgi’ were combined into one ‘Endosome-Lysosome-Golgi’ category.

At first, interactomes were aggregated across CI, WT, CA paralog states. Next, over-representation analysis using g:profiler2’s ‘gost’ (version e113_eg59_p19_f6a03c19) functions was conducted for the subcellular interactome of each core protein. Over-represented terms were retrieved for relevant organelles (‘plasma_membrane’, Endosome-Lysosome-Golgi, ‘actin_cytoskeleton’, ‘ER’, ‘nucleus’, and ‘unclassified’). In order to select the most relevant GO:BP and Reactome terms, the subcellular stratified interactomes were uploaded and analyzed using STRING-DB. Enriched terms explaining the highest number of interactors were manually selected for the final dot plot.

#### DISGENET enrichment

Only the CA network was considered for disease enrichment. Tier overlapping preys (e.g preys shared in RAS-RAF, RAF-MEK,etc.) were used as input for disease enrichment using the disgenet2r library (DISGENET v25.2). For the analysis of CA interactomes, bait proteins were aggregated into one entity per state, in the case of a protein having two activating mutants (e.g the interactomes of BRAF G469A and BRAF V600E were aggregated into CA-BRAF).

#### Cancer-Gene-Census (CGC) analysis

The CGC data set was downloaded in September 2025. We restricted the analysis to Tier 1 genes, representing the most clinically relevant and highest confidence cancer drivers, and excluded genes annotated as “fusion” in the “Role in cancer” column, in order to focus on individual gene effects. The interactome was filtered by the remaining genes from the filtered CGC table.

#### Network data

The following network resources were used:

Intact^73^: Release 251 - September 2025 - downloaded on September 10th 2025.

SIGNOR^33^: Jul2025_release.txt downloaded on 15th August 2025.

CORUM^47^: Corum 5.1 release (2025-01-21) downloaded on 31st July 2025.

STRING-DB^74^: STRING_v12.0 fetched from STRING API in September 2025.

#### Network visualization

Network layouts for the huMAP3.0 complexes, CA network, and DisGeNET diseases were constructed in Python using networkx. At first, the bait proteins were placed in an outside ring using the circular_layout function. The prey nodes were positioned inside using a spring_layout which positions were manually refined in Cytoscape. In addition, for the CA network edge bundling was performed in CytoScape, using default parameters. Networks using the STRING-DB resource were retrieved using the STRING-DB API (https://www.string-db.org/api/json/network?identifiers=<ID_1>%250d<ID_2>%250d&<ID_N>species=9606&required_score=400&network_type=<physical|functional>&version=v12.0). Physical edges were filtered at a minimum STRING score of 400, functional edges at a higher minimum score threshold of 750, to retrieve only high-confidence functional associations, similar as in Barrio-Hernandez et al.^85^

#### Figure generation and general code availability

Data analysis for this study was conducted using the R and Python programming languages. Figures were plotted in R using the ggplot, igraph, RColorBrewer, and cowplot libraries. Plotting and analysis functions from the amica web platform were used and extended. All analysis and visualization code will be available on github and archived on Zenodo after publication.

### QUANTIFICATION AND STATISTICAL ANALYSIS

The statistical approach for each experiment is described in the corresponding STAR Methods section and or in the figure legends.

---

**Supplementary Figure 1.**
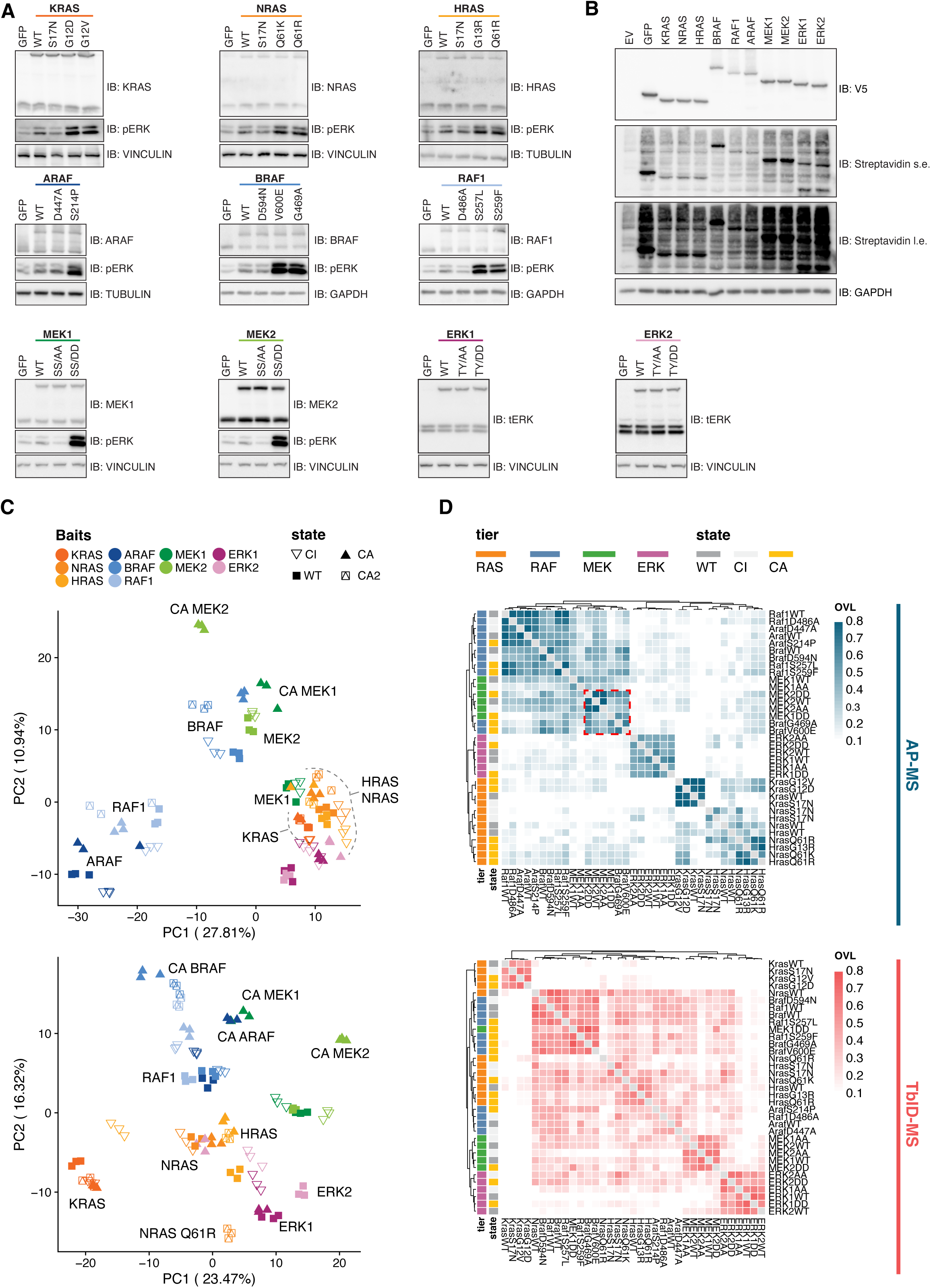
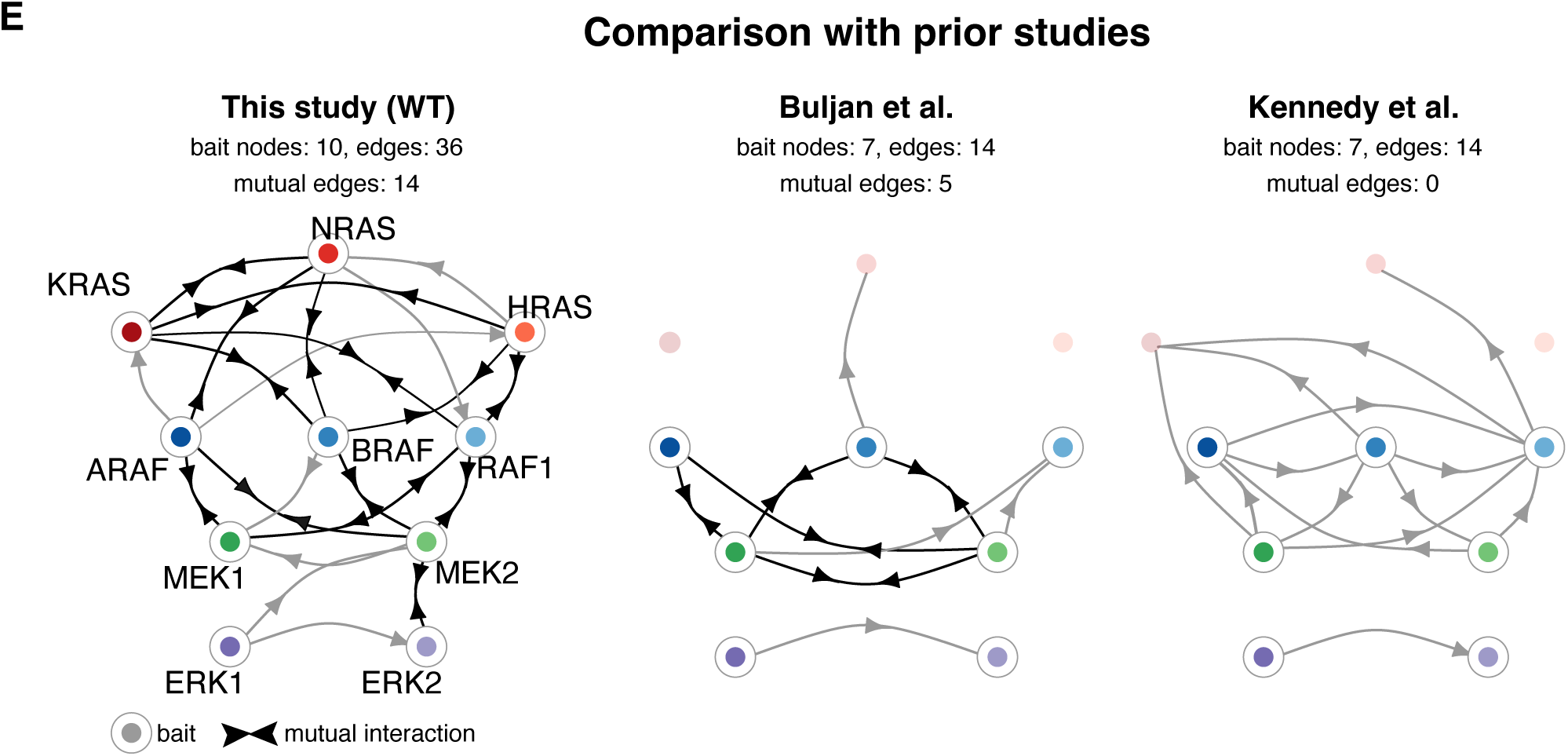
Quality controls and benchmarks. (A) Immunoblots showing bait expression at near endogenous levels in the respective cell lines as assessed by antibodies specific for the proteins of interest. The GFP-TbID-V5 cell line is shown as a control. (B) Similar proximity labeling efficiency across baits as assessed by V5 and streptavidin immunoblotting. Endogenous ERK phosphorylation is used as a readout of CA mutant activity. In A and B, GAPDH, TUBULIN or VINCULIN were used as loading controls. EV = empty vector; s.e. = short exposure; l.e. = long exposure. (C) Principle component analysis (PCA) on bait - prey intensity matrices. Triplicate WT, CI or CA samples are color-coded. States are represented by different shapes. (D) Hierarchical clustering of pairwise overlap coefficients (OVL) between individual ERK pathway interactomes. HCPPIs are color-coded by state and tier. The same color-code is maintained throughout all figures. (E) Comparison of directed networks of core pathway HCPPIs constructed from the data in this study and in the two largest IntAct MS datasets using ERK pathway core proteins as baits. The relationship between baits and preys is indicated by the direction of the arrowheads. Buljan et al. and Kennedy et al. did not use RAS baits.

**Supplementary Figure 2.**
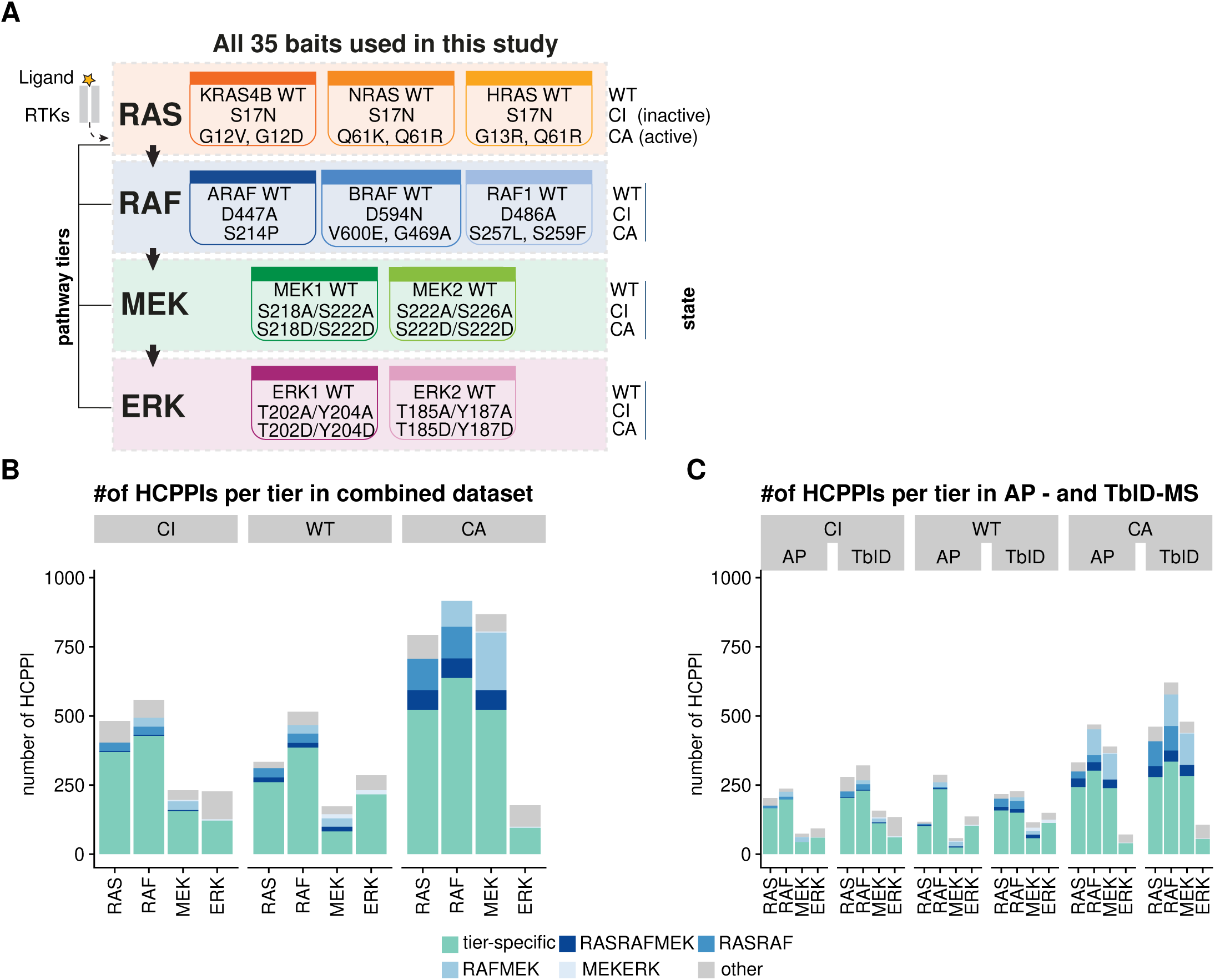
Tier overlap increases in the CA network. (A) RAS/RAF/MEK/ERK pathway. Each of the 10 core proteins is represented by a different color and is mapped in 3 states (WT, CI, and CA), resulting in the 35 bait proteins depicted. The same color code is maintained throughout all figures. (B, C) Bar plots showing the number of HCPPIs in each of the four tiers in the CI, WT, and CA states. Tier-specific HCPPIs as well as HCPPI found in contiguous are indicated. Other = preys shared by non-contiguous tiers. (B), AP-MS and TbID-MS datasets; (C) combined dataset

**Supplementary Figure 3.**
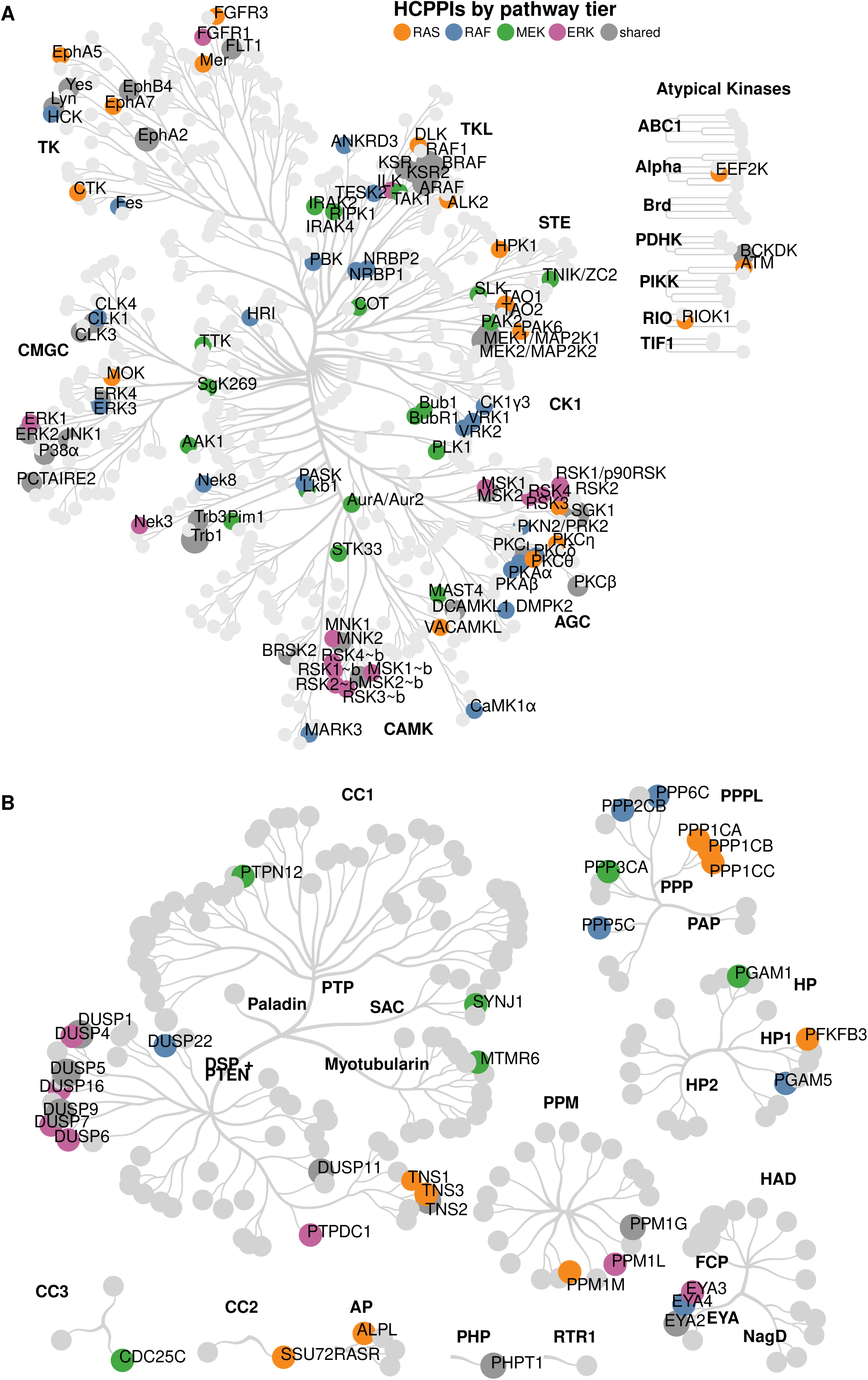
Pathway interactions with kinases and phosphatases. (A, B) AP-MS/TbID-MS interactomes mapped on the kinome/phosphatome evolutionary tree. HCPPIs are color-coded by tier and aggregated across states. Kinase/phosphatase families are in bold. Shared = proteins interacting with multiple tiers. All HCPPIs mapping to the trees are labeled.

**Supplementary Figure 4.**
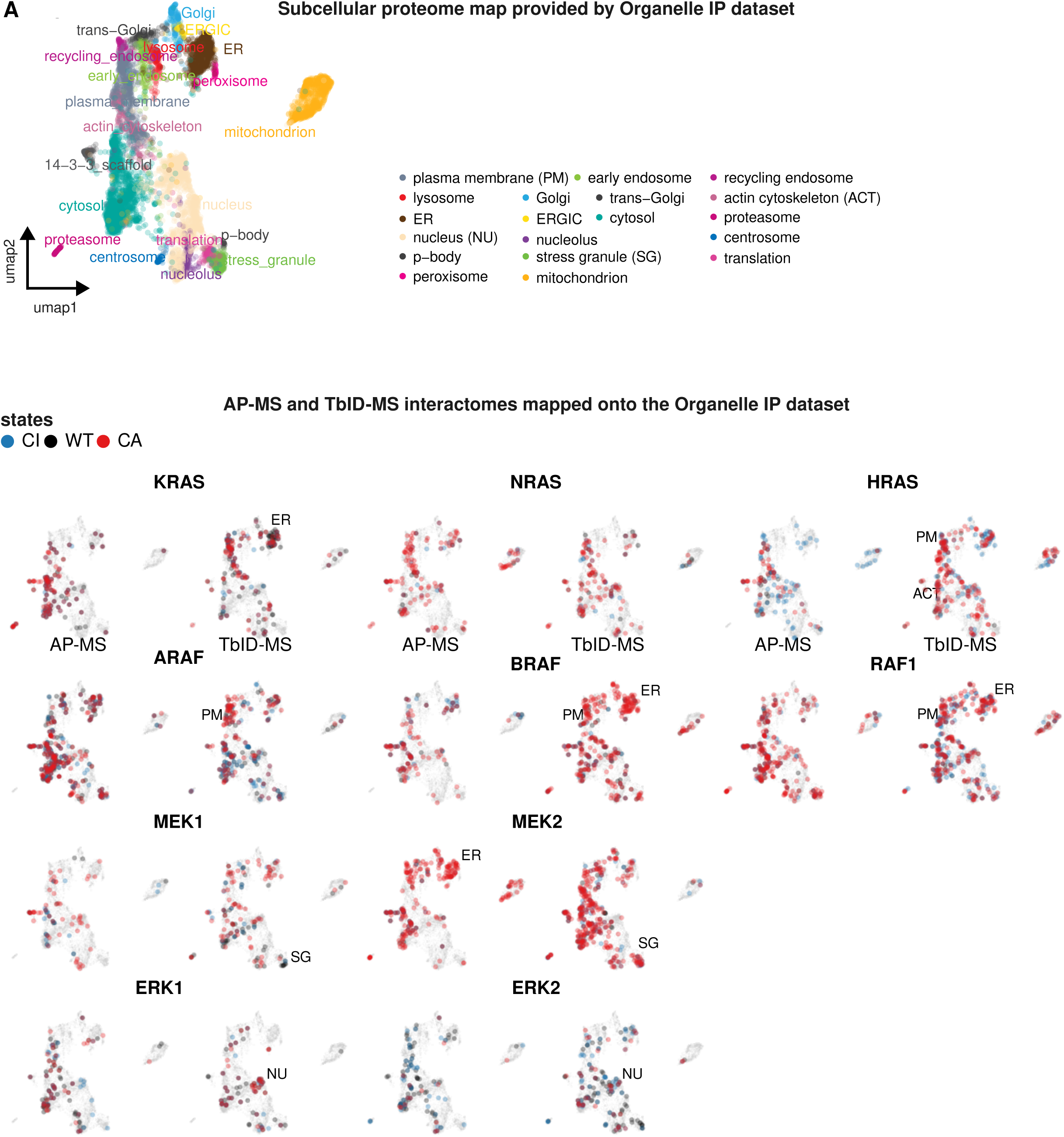
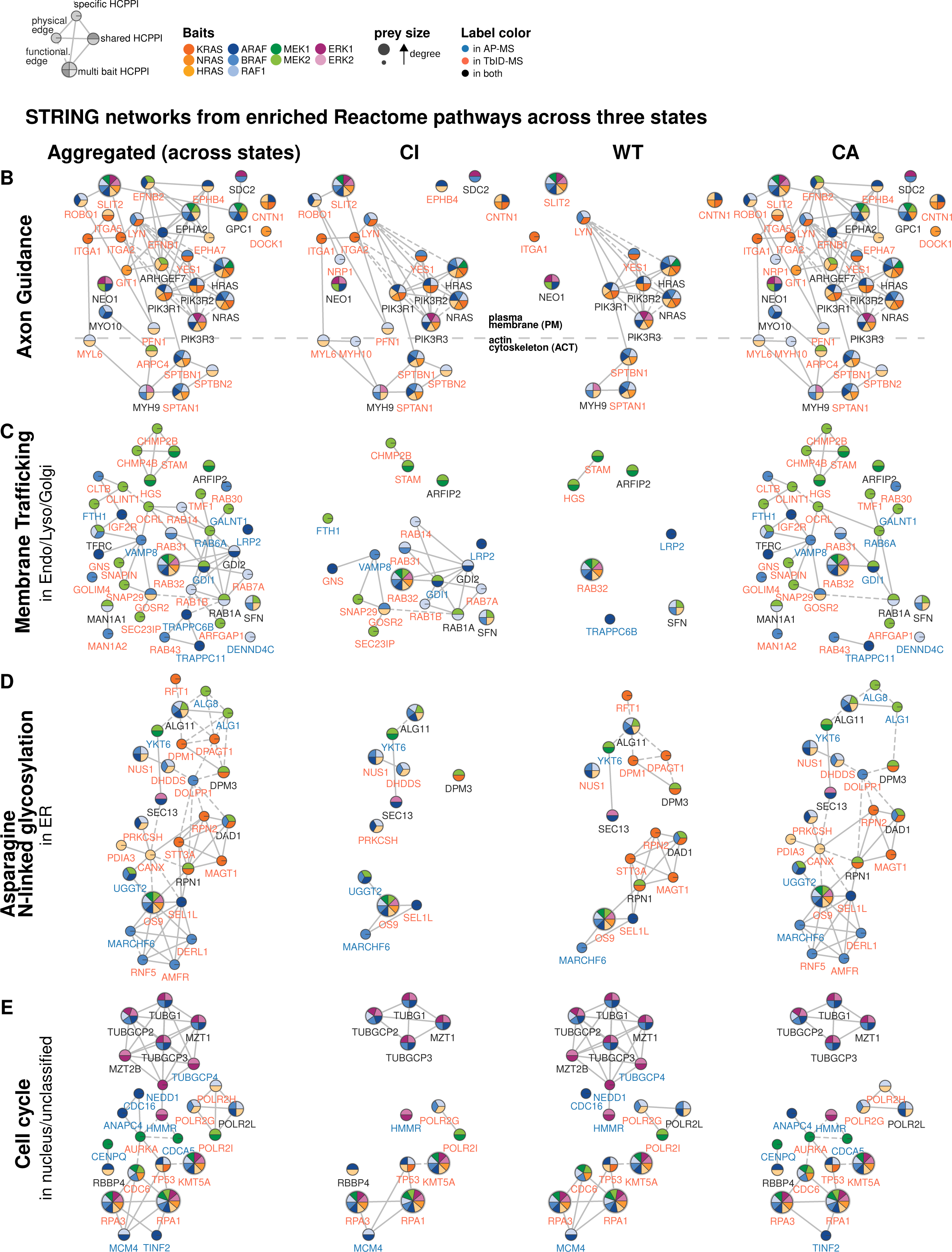
AP-MS and TbID-MS interactomes mapped onto the Organelle IP dataset. (A) Two-dimensional UMAPs showing the HCPPI identified in this study mapped onto the 19 subcellular compartments annotated by the <u>organelleIP</u> dataset. Each dot represents an individual HCPPI and it is color-coded according to the state (CI, WT, CA) of its bait. HCPPIs detected by AP-MS are shown on the left, TbID-MS on the right. The original subcellular UMAP is shown at the top as reference. PM = plasma membrane; ACT = Actin cytoskeleton; ER = endoplasmatic reticulum; SG = stress granules; NU=nucleus. (B-E) State-specific STRING networks of HCPPIs involved in selected Reactome pathways. Nodes are colored according to the paralogs they interact with. Node size is proportional to the number of interactions with the baits (node degree). Blue labels = nodes detected by AP-MS; red = detected by TbID; black = detected by both. Solid edges = STRING physical interactions (minimum score 0.4); dashed edges = STRING high-confidence functional interactions (minimum score 0.75).

**Supplementary Figure 5.**
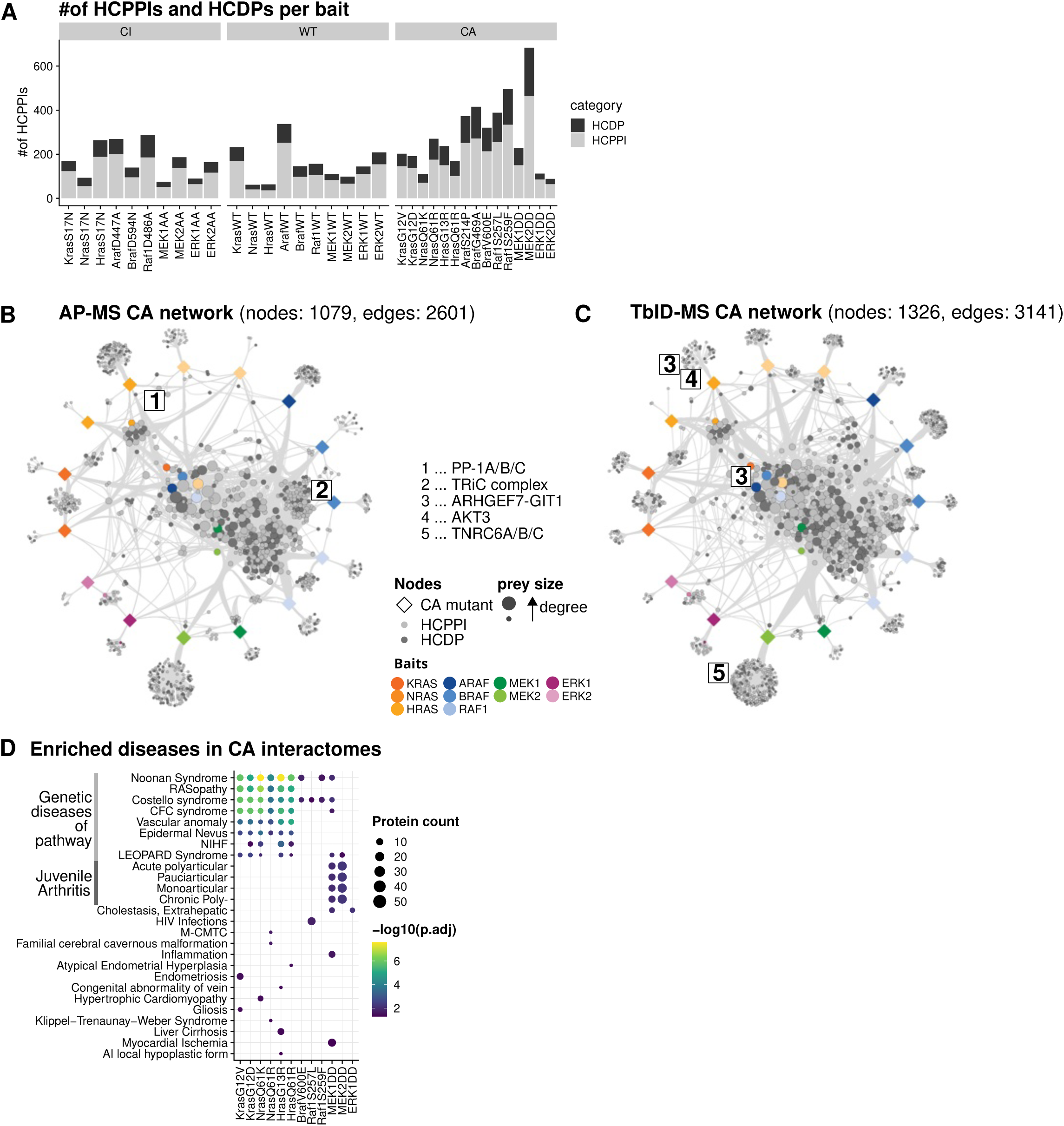
Pathway involvement in disease. (A) Bar plot showing the number of HCDPs/HCPPIs per bait in the CI, WT, and CA states. The number of HCDPs increased in the CA network. (B-C) Network representation of CA interactomes. HCDPs (DISGENET score > 0.66) are in dark grey; HCPPIs representing endogenous pathway proteins are color-coded. HCPPI node size is proportional to the number of interactions it has with the baits. Edges with similar start and end nodes were bundled in Cytoscape using default parameters to emphasize network topology. Selected proteins unique to one of the methods are highlighted. (D) DISGENET enrichment analysis of combined AP-MS and TbID-MS interactomes of individual CA baits. Dot size is proportional to the number of proteins in each DISGENET term; color scale = -log10 (adj. p-value).

**Supplementary Figure 6.**
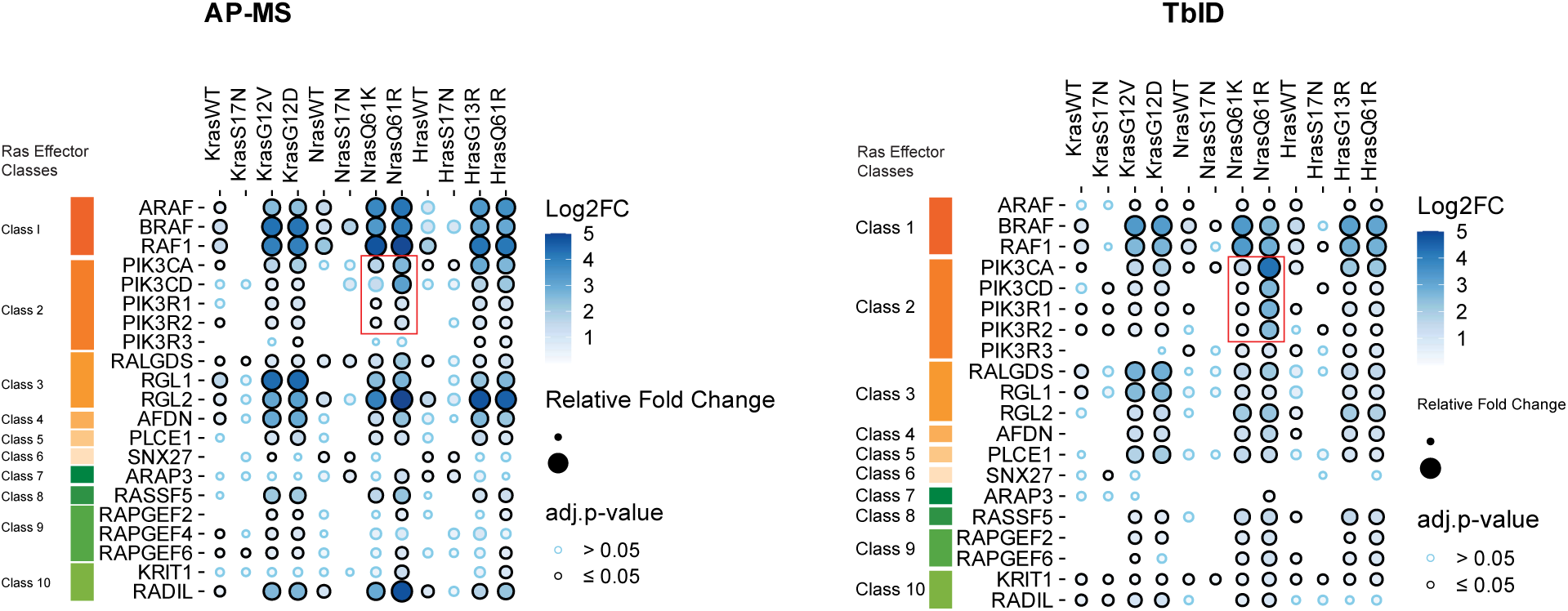
RAS effectors identified in AP-MS and TbID-MS. Dot plots show a curated list of detected RAS effectors. Visualized as in Figure 6A.

**Supplementary Figure 7.**
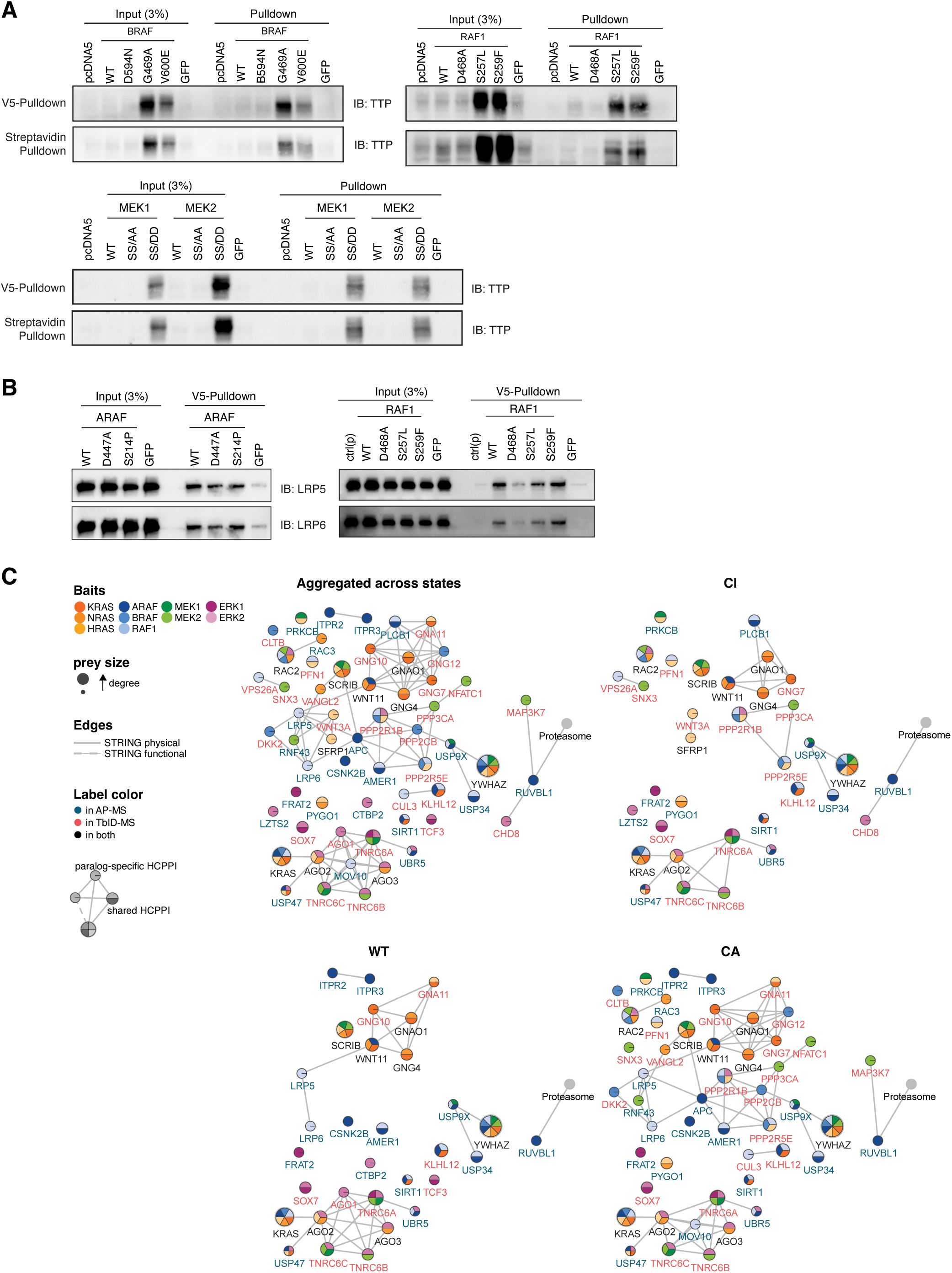
Cross-talk with mRNA metabolism and WNT. (A-B) Immunoblot validation of the interaction of ZFP36/TTP (A), LRP5 and LRP6 (B) with RAF and MEK by V5 and/or streptavidin pulldown. (C) State-specific STRING networks of HCPPIs involved in selected Reactome pathways. Nodes are colored according to the paralogs they interact with. Node size is proportional to the number of interactions with the baits (node degree). Blue labels = nodes detected by AP-MS; red = detected by TbID; black = detected by both. Edges represent STRING physical interactions (minimum score 0.4).

## Notes

### Competing Interest Statement

The authors have declared no competing interest.

